# Beyond autoantibodies: Biological roles of human autoreactive B cells in rheumatoid arthritis revealed by whole transcriptome profiling

**DOI:** 10.1101/144121

**Authors:** Ankit Mahendra, Xingyu Yang, Shaza Abnouf, Daechan Park, Sanam Soomro, Jay RT Adolacion, Jason Roszik, Cristian Coarfa, Gabrielle Romain, Keith Wanzeck, S. Louis Bridges, Amita Aggarwal, Peng Qiu, Sandeep Krishna Agarwal, Chandra Mohan, Navin Varadarajan

## Abstract

Although the contribution of B-cell derived autoreactive antibodies to rheumatoid arthritis (RA) has been studied extensively, the autoantibody-independent roles of B cells in the progression of the disease is not well-defined. Here we present the first comprehensive transcriptome profile of human autoreactive B cells in an autoimmune disease by performing RNA-sequencing of citrulline-specific B cells from RA patients. In order to facilitate a comprehensive understanding of the profile of these citrulline-specific (RA-CCP^POS^) B cells, we performed comparative analyses to both citrulline-negative (RA-CCP^NEG^) B cells from the same donors, and identified 431 differentially expressed genes (DEGs); and hemagglutinin-specific (HA) B cells from healthy individuals and identified 1658 DEGs. Three-way comparisons of these B cell populations demonstrated that RA-CCP^POS^ B cells, in comparison to the RA-CCP^NEG^ B cells, demonstrate a potential role in protein citrullination and inflammation; RA-CCP^POS^ B cells in comparison to HA-specific B cells demonstrate RA-specific signatures like the expression of pro-inflammatory cytokines, chemokines, costimulatory molecules and B-cell activation cascades; and all B cells from RA patients demonstrated a significant impact of the multitude of TNF signaling pathways. Furthermore, transcription factor profiling suggested that cyclic AMP (cAMP) related pathways and downstream signaling molecules are selectively enriched in RA-CCP^POS^ cells in comparison to the other two B cell subsets. We advanced the understanding of the citrulline reactive B cells in RA pathophysiology by documenting and validating two novel observations in independent cohorts of patients: (1) the expression of IL15Rα is restricted to citrulline-specific cells within RA patients and the concentration of soluble IL15Rα is elevated in the sera of RA patients, (2) B cells from RA patients are capable of producing epidermal growth factor ligand, amphiregulin (AREG) which in turn has a direct impact on the mechanistic effectors of RA, osteoclasts and fibroblastlike synoviocytes (FLS). Overall, our comprehensive dataset identifies several existing FDA-approved drugs that can potentially be repurposed for RA and can serve as a foundation for studying the multi-faceted roles of B cells in other autoimmune diseases.

## Introduction

RA is an autoimmune disease characterized by swelling of the synovial membrane with the robust infiltration of cells of both the innate and adaptive immune system: B cells, T cells, macrophages, dendritic cells, neutrophils and mast cells^1^. The identification of citrullination, the post-translational modification of arginine residues to citrulline in proteins, catalyzed by the enzymes peptidyl arginine deaminases (PADs), as one of the key factors mediating the breach in tolerance and eliciting anti-citrullinated protein antibody (ACPA) responses, has been a major milestone for RA^2,3^. The appearance of ACPA in circulation precedes the onset of clinical disease, and ACPA positivity has a sensitivity of 60-70%, and a specificity of greater than 90%, for the diagnosis of RA^2^. ACPA have shown to trigger human immune effector functions including activation of the complement system and the ability to engage activating Fcγ receptors^4^. Lastly, it has been demonstrated that ACPA can catalyze bone erosion commonly seen in RA patients^5^.

Although the indispensable role of B cells in autoantibody production is well recognized, their autoantibody-independent contributions are not as well-defined. Systemic depletion of B cells using rituximab that targets B cells expressing human CD20, a phenotypic cell surface marker, has been shown to be effective clinically for the treatment of a subset of RA patients^6^. While studies have supported the idea that patients with higher autoantibody titers are likely to respond to rituximab^7,8^, the clinical efficacy of rituximab treatment is not necessarily correlated with a decrease in autoantibody titers^9^. This, in turn, implies a more expanded role for B cells in autoimmune pathogenicity beyond antibody production. Preclinical and clinical data have suggested many different functions of B cells in RA including a role as antigen presenting cells (APCs) that can process and present peptides enabling T-cell activation and proliferation^10,11^; direct secretion of pro-inflammatory cytokines like tumor necrosis factor (TNF) and interleukin-6 (IL-6)^12,13,14^; supporting the organization of tertiary lymphoid tissues within the inflamed synovium^10,15,16^; and impacting bone homeostasis through the secretion of receptor activator of nuclear factor kappa-B ligand (RANKL)^17,18^. Despite these efforts, a direct interrogation of the roles of the autoreactive B cell compartment, and how they differ from other B cells from the same donor or from B cells from healthy individuals, has not been accomplished.

One of the major challenges in profiling autoreactive B cells within humans is the low frequency of these cells in peripheral blood (< 0.1 % of all B cells in circulation)^19^. We developed and validated a flow-cytometry based assay for the reliable detection of cyclic citrullinated peptide (CCP) specific B cells *ex vivo*. We then performed RNA-sequencing (RNA-seq) on these small numbers of cells and compared the whole transcriptome profile of CCP-specific B cells (RA-CCP^POS^) with that of CCP-negative B cells (RA-CCP^NEG^) from the same donor and hemagglutinin-specific (HA^POS^) B cells from healthy individuals elicited upon vaccination with seasonal influenza virus. Our data identifies a novel role of RA-CCP^POS^ B cells as an activator of the epidermal growth factor (EGF) pathways by secretion of EGF receptor ligand amphiregulin (AREG) under pro-inflammatory conditions, and also identifies IL15Rα, as a specific biomarker of RA-CCP^POS^ B cells. Overall, our data suggest that besides being a source of autoantibodies, B cells may also play direct roles in the inflammatory cascades and osteoclastogenesis in RA. Significantly, while some of the identified genes and pathways are the targets of approved therapies for RA, our data also illustrates that drugs approved for other indications might be useful in targeting the RA-CCP^POS^ B cell compartment in RA. More broadly, to the best of our knowledge, this is the first comprehensive study on the transcriptome profile of human RA-CCP^POS^ B cells in autoimmune disease and will serve as a resource to further investigate the role of B cells autoimmunity.

## Results

### Flow-sorting of antigen-specific B cells

We developed a dual-labeling, flow sorting method using both cyclic citrullinated (CCP) and cyclic arginine peptides (CAP) to isolate RA-CCP^POS^ B cells. In order to verify the purity of our sorting method, an equal number of cells within the CCP^POS^AP^NEG^ (hereafter referred to as RA-CCP^POS^ B cells), CCP^NEG^AP^POS^ and CCP^NEG^AP^NEG^ (hereafter referred to as RA-CCP^NEG^) populations (**Figure 1A**) were sorted in 96 well plates and grown *in vitro* for 14 days as described previously^19^. The purity of our sorting strategy was validated by testing the supernatants after *in vitro* culture, which confirmed that only the immunoglobulins secreted in B-cell culture established from the RA-CCP^POS^ B cell population demonstrated a specific reactivity towards the CCP (**Figure 1B-C**). After validation of our sorting strategy, a total of 350-1000 RA-CCP^POS^ B cells (0.01 – 0.1 %) from the blood of four RA patients were used directly for the preparation of cDNA libraries *ex vivo* to ensure minimal perturbations to the transcriptional profile (**Table 1**). Both RA-CCP^POS^ and RA-CCP^NEG^ B cells were confirmed to be predominantly of the memory phenotype based on the surface expression of CD27 (**Figure S.1**).

**Figure 1.**
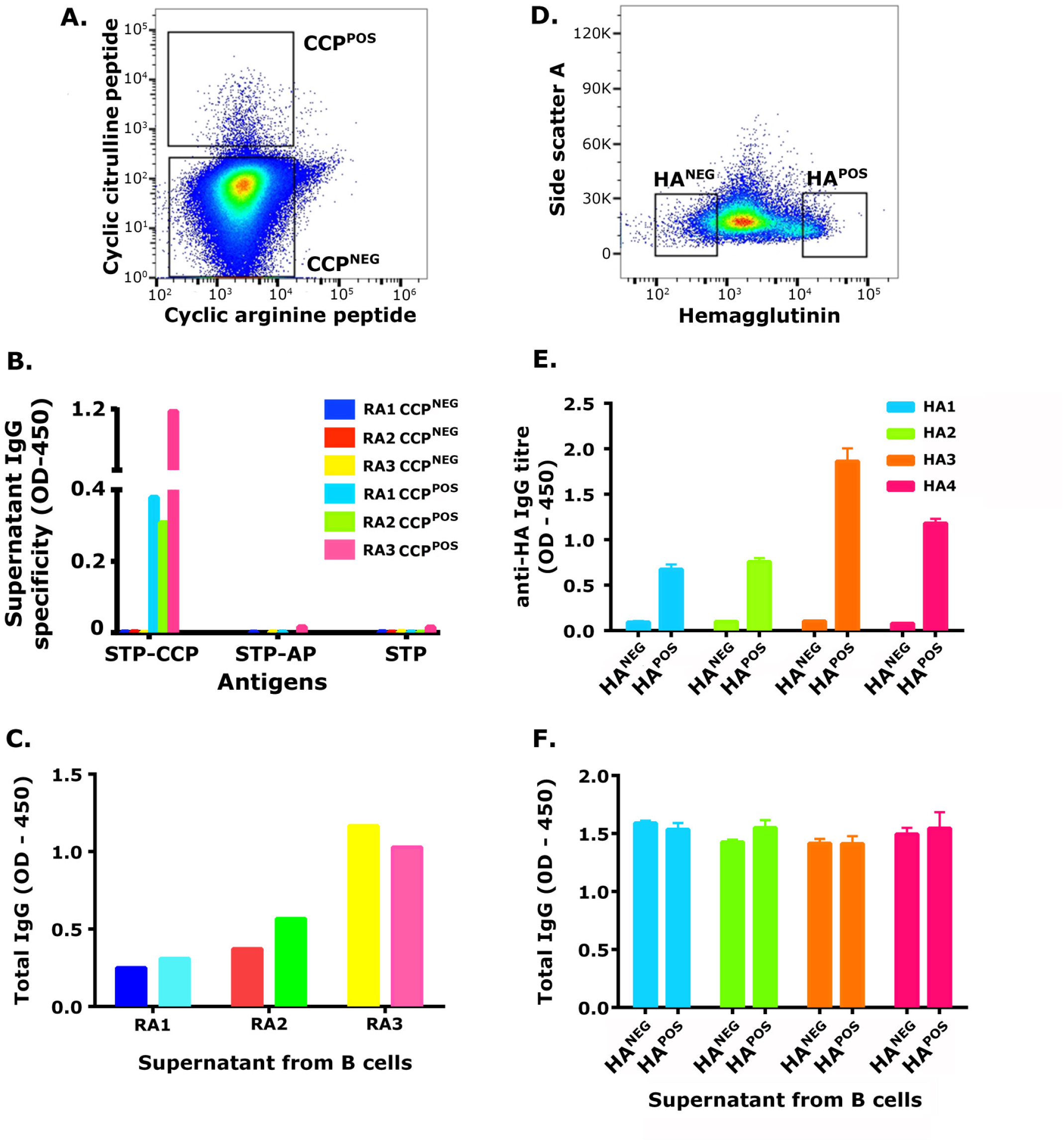
Isolation of an enriched population of RA-CCP^POS^ and HA-specific B cells. **A**. RA-CCP^POS^ B cells were flow sorted using CCP-biotin-streptavidin tetramers and gated as CCP^POS^AP^NEG^, ELISA on supernatants testing for (B) antigen reactivity, and (C) total Ig, from B cells expanded and differentiated *in vitro* (n = 3). **D**. Representative flow plot displaying gating strategy for sorting HA^POS^ B cells. **E**. ELISA on supernatants testing for (E) HA reactivity and (F) total Ig from control and HA-specific B-cell populations (n = 4).

**Table 1.**
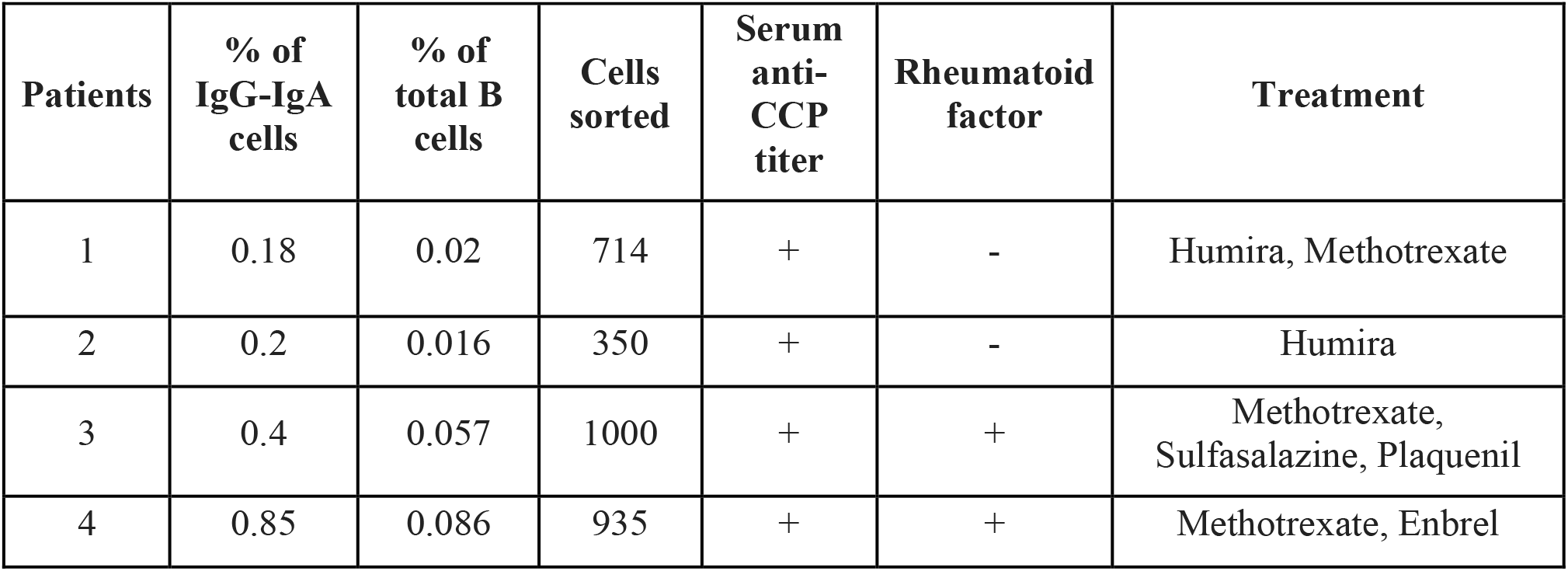
Patient characteristics of RA donors.

In order to have a comparative analysis of B-cell transcriptome profile during autoimmunity versus normal immune response to a pathogen, HA-specific B cells (hereafter referred to as HA^POS^ B cells) were isolated from blood of four healthy individuals vaccinated with the seasonal influenza vaccine. Our ability to enrich for HA^POS^ B cells was validated by the same three step procedure used for RA-CCP^POS^ B cells: (a) antigen labeling and flow-sorting a total of 3500 HA^POS^ and HA^NEG^ cells from the PBMCs of these vaccinated donors, (b) *in vitro* expansion and differentiation, and (c) ELISA testing for HA-reactivity on the culture supernatants (**Figure 1D-F**). Subsequent to validation, 1000 HA^POS^ B cells from the same four donors were used to construct cDNA libraries *ex vivo* for RNA-sequencing (RNA-seq).

### Transcriptome analysis revealed that RA-CCP^POS^, RA-CCP^NEG^, and HA^POS^ B cells could be distinguished based on the differentially expressed genes

The cDNA libraries generated *ex vivo* from 12 samples (4 paired RA-CCP^POS^ B cells and RA-CCP^NEG^ populations, and 4 HA^POS^ B cell populations) were barcoded, pooled and sequenced using 76bp paired-ends to yield a minimum of 17 million reads per library. After validation of the RNA-seq populations (**Figure S.2**), differential analyses using the DEseq package revealed that 1658 genes (false discovery rate, FDR < 0.1) were differentially expressed in RA-CCP^POS^ B cells in comparison to the HA^POS^ B cells, and 431 genes were identified as differentially expressed genes (DEGs) in comparison to the RA-CCP^NEG^ B cells (**Table S.1** and **S.2**). We utilized t-distributed stochastic neighbor embedding (t-SNE), to demonstrate that these identified DEGs could clearly resolve the three distinct cellular populations^20^ (**Figure 2A**). As expected, both the number of identified DEGs and their relative change in expression was lower in comparing the RA-CCP^POS^ and RA-CCP^NEG^ B cells from the same donors, in contrast to comparing the RA-CCP^POS^ B cells to the HA^POS^ B cells (**Table S.1** and **S.2**), with only 22 DEGs being conserved in both comparisons.

**Figure 2.**
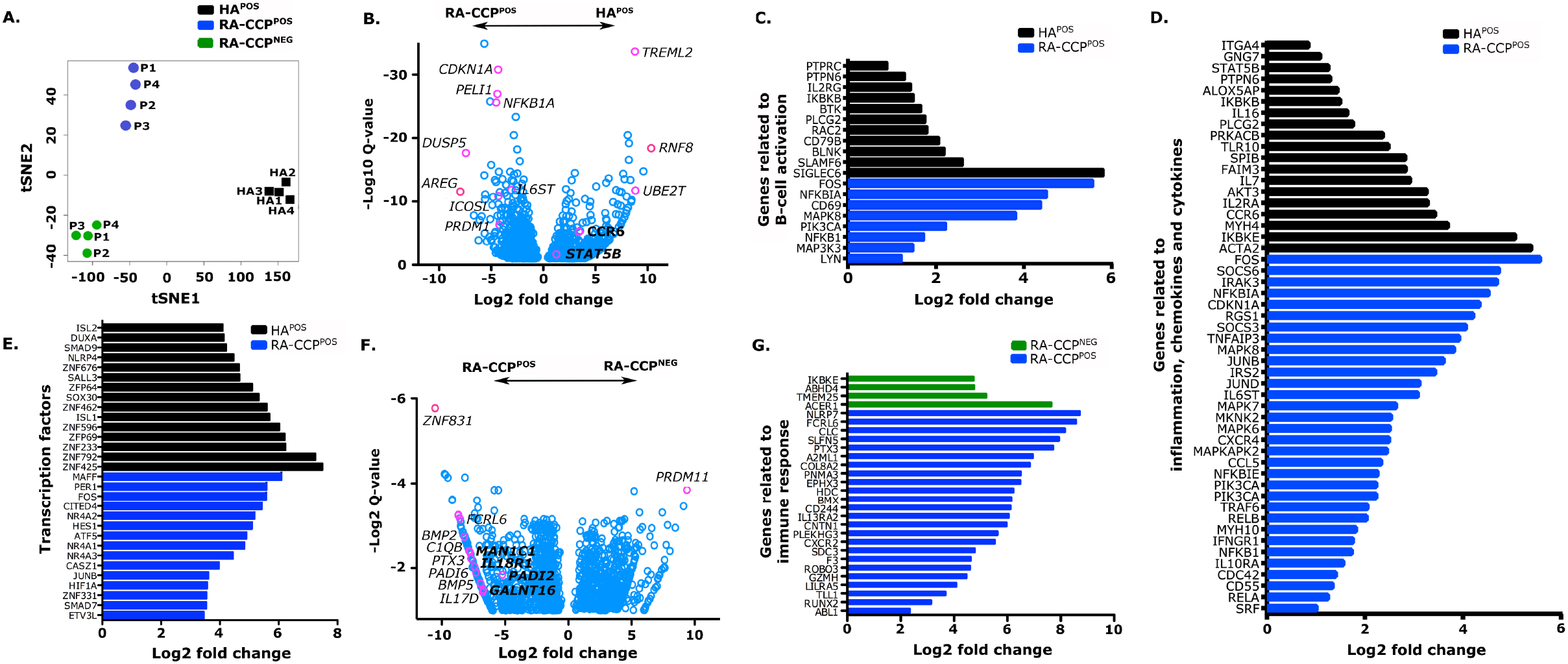
RA-CCP^POS^, RA-CCP^NEG^, and HA^POS^ B cells can be differentiated based on DEGs. **A**. t-Distributed Stochastic Neighbor Embedding (t-SNE) visualization of sample relationships using the differentially expressed genes. **B**. Volcano plots generated using fold change and *q-value* of DEGs shows enrichment of genes related autoimmunity and inflammation in RA-CCP^POS^ B cells in comparison to HA^POS^ B cells. (**C-E**). Gene ontology plots indicating fold change in expression of genes related to B-cell activation, inflammation, chemokines and cytokines, and top fifteen transcription factors that are differentially enriched in RA-CCP^POS^ versus HA^POS^ B cells. **F**. Volcano plot generated using fold change and *q-value* of DEGs between RA-CCP^POS^ B cells and RA-CCP^NEG^ B cells from the same donors. **G**. Gene ontology plot depicting fold change in expression of genes related to immune responses that are highly expressed in RA-CCP^POS^ B cells in comparison to RA-CCP^NEG^ B cells from the same donors.

### Comparisons between RA-CCP^POS^ and HA^POS^ B cells

A number of candidate DEGs that are well validated in autoimmunity including the cyclin kinase p21 (*CDKN1A*)^21^, ubiquitin ligase *PelI1*^22^, and the costimulatory molecule *ICOSLG*^23^ were identified. The epidermal growth factor ligand, amphiregulin (*AREG*) was identified as the transcript with the largest change in expression (**Figure 2B**). By classifying these same DEGs into the Gene Ontology (GO) categories, we grouped them as being related to B-cell activation, genes related to inflammation, and transcription factors. We observed differences in the expression of genes related to B-cell activation (**Figure 2C**) with increased expression of kinases like phosphatidylinositol-4,5-bisphosphate 3-kinase catalytic subunit alpha (*PI3KCA*)^24^ and mitogen-activated protein kinases (*MAPK7/MAPK8*) known to cause autoimmunity under deregulated activation^25,26,27^. In addition, several genes related to inflammation including signaling molecules like *TNFAIP3*^28^ and *IL6ST*^29^, and T-cell recruiting chemokine-like *CCL5*^30^ and chemokine receptor *CXCR4*^31^ were upregulated in RA-CCP^POS^ B cells (**Figure 2D**).

Analysis of the differentially expressed transcription factors (TFs) revealed upregulation of positive regulatory domain I-binding factor 1 (*PRDM1*) and ETS variant 3 (*ETV3* and *ETV3L*) within RA-CCP^POS^ B cells; these TFs are known to inhibit c-Myc and consequently cell growth^32^, while at the same time promoting B-cell differentiation and immunoglobulin (Ig) secretion^33^. Similarly, the small MAF family TFs (*MAFF* and *MAFG*) that can directly impact Ig secretion were enriched in these RA-CCP^POS^ B cells^34^. Three separate members of the NR4A orphan nuclear receptors (*NR4A1, NR4A2*, and *NR4A3*), whose functions in B cells are not well-defined, but roles as master transcription factors that can regulate cell fate by influencing metabolism, proliferation, and apoptosis within cells of the hematopoietic lineage is well-documented^35,36,37^, were also upregulated in RA-CCP^POS^ B cells (**Figure 2E**).

We also sought to determine if there was a B-cell specific change in expression of genes known to be high-risk loci in RA, documented through large-scale genetic efforts^38,39^. Out of the known RA-specific loci, 9 genes were also identified as DEGs (*SH2B3, CCR6, ILF3, TXNDC11, PTPRC, PRDM1, TNFAIP3, TRAF6*, and *LBH*) [**Figure S.3**].

### Comparisons between RA-CCP^POS^ and RA-CCP^NEG^ B cells

The candidate DEGs upregulated within the RA-CCP^POS^ population in comparison to RA-CCP^NEG^ B cells within these same donors included: pentraxin *PTX3* that recognizes pathogen-associated molecular patterns (PAMP) and its binding partner in the complement cascade (*C1QB*)^40,41^; the bone morphogenetic proteins (*BMP2* and *BMP5*)^42^; and the peptidyl arginine deaminase citrullinating enzymes (*PADI2* and *PADI6*)^43^ [**Figure 2F**]. Additionally, several immune-related transcripts like the inflammasome-associated protein *NLRP7^44^* and the Fc receptor-like protein *FCRL6^45^* were also upregulated within RA-CCP^POS^ B cells (**Figure 2D**).

### Cytokine and signaling pathways enriched in RA-CCP^POS^ B cells

The differentially expressed pathways were obtained by matching the expression dataset using Ingenuity pathway analysis (IPA), filtering the pathways directly related to immune cells and their functions, and visualizing as a heatmap by using the log *p-values* (**Figure 3A**). In order to facilitate comparisons, we identified three groups of pathways that were enriched: both in the RA-CCP^POS^ and RA-CCP^NEG^ B-cell populations in comparison to the HA^POS^ B cells (group 1), only in the RA-CCP^POS^ population but not the RA-CCP^NEG^ or HA^POS^ B cells (group 2), and only in the RA-CCP^POS^ population in comparison to the RA-CCP^NEG^ B cells (group 3) [**Figure 3A**]. The group 1 pathways comprised of B-cell activation pathways (B-cell receptor signaling, PI3K/Akt, and p38 MAPK), toll-like receptor-based activation (TLR signaling and LPS-stimulated MAPK pathways), and the TNF superfamily ligand-mediated signaling (CD40, APRIL signaling) and their receptor signaling pathways (TNFR1, TNFR2, CD27, RANK, and CD40). The group 2 pathways comprised of inflammatory cytokine signaling (IL6, IL8, and IL-17A) and RA-specific signatures (role of macrophages, endothelial cells and osteoclasts in rheumatoid arthritis); B-cell activation/homeostasis (PTEN, NF-κB, and NFAT activation) and function (phagosome maturation, FcγRIIb signaling, and antigen presentation). Finally, group 3 showed enrichment in the pathways for protein citrullination, G-protein coupled receptor signaling (Gαq signaling) and cyclic AMP (cAMP) signaling (**Figure 3A**). Overall, these data suggest that while TNF signaling has a global impact on all B cells from RA patients (group 1), the RA-CCP^POS^ B cells in comparison to the RA-CCP^NEG^ B cells demonstrate a potential role in protein citrullination and effector functionality.

**Figure 3.**
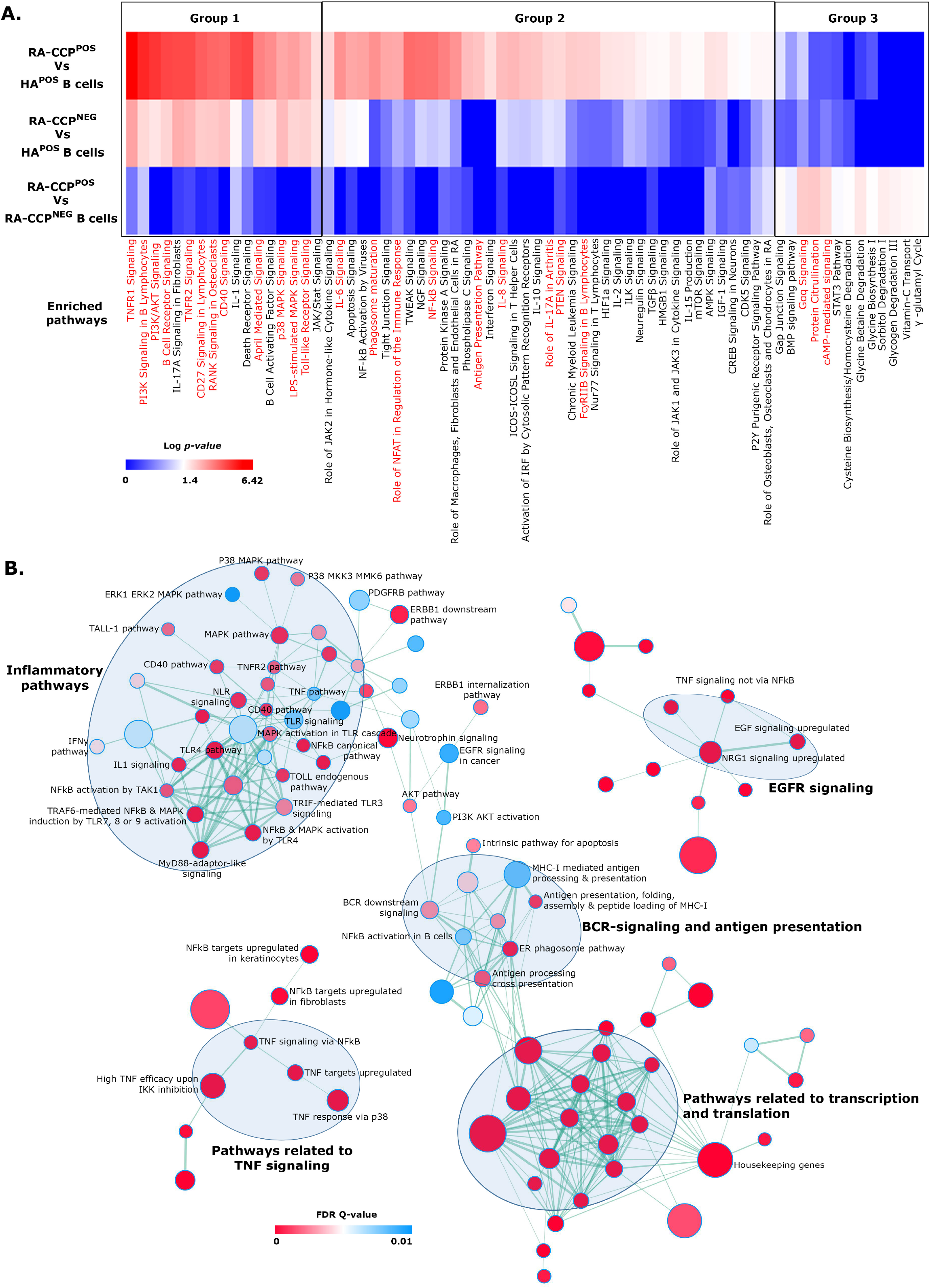
Enrichment maps of pathways enriched in RA-CCP^POS^ B cells as compared to HA^POS^ B cells. **A**. Heat map generated using log-transformed *p-values* of DE-pathways from IPA indicated enrichment of pro-inflammatory pathways in RA-CCP^POS^ B cells as compared to HA^POS^ B cells and upregulation of protein citrullination in RA-CCP^POS^ B cells versus RA-CCP^NEG^ B cells. Highlighted pathways are marked in red. **B**. The GSEA-derived enriched C2 curated pathways in RA-CCP^POS^ B cells were plotted using the enrichment map application in Cytoscape using a cutoff FDR *q-value* = 0.01 and a *p-value* = 0.005. Nodes (circles colored red and blue) represent pathways and the edges (green lines) represents overlapping gene among pathways. The size of nodes represents the number of genes enriched within the pathway and the thickness of edges represents the number of overlapping genes. The color of nodes was adjusted to an FDR *q value* range from 0 - 0.01. Clusters of pathways are labeled as groups with a similar theme. All pathways represented here are enriched in RA-CCP^POS^ B cell populations.

Based on the substantially larger number of pathways and greater changes in expression, identified in the RA-CCP^POS^ B cells in comparison to the HA^POS^ B cells by IPA (groups 1 and 2 combined), we performed gene-set enrichment analyses (GSEA) comparing these two populations^46^. We interrogated the changes in these populations against the Molecular Signatures Database (Hallmark and C2 curated gene sets). As shown in **Figure 3B**, five major clusters of pathways were significantly upregulated (FDR *q-value* < 0.1) in the RA-CCP^POS^ B cells: transcription and translation; B-cell receptor signaling; EGFR signaling; TNF signaling; and inflammatory cytokines and chemokines.

### IL5RA expression is restricted to RA-CCP^POS^ B cells within RA patients

As outlined above, multiple pathways that share genes with the TNF pathway and genes that are targets of TNF were enriched in RA-CCP^POS^ B cells from RA patients (**Figure 4A**). Similarly, the inflammatory response and the IL-6-Jak-STAT3 pathways were also upregulated in RA-CCP^POS^ B cells in comparison to HA^POS^ B cells (**Figure 4B**), and the IL-6 signal transducer (*IL6ST*) was also a DEG with elevated expression within RA-CCP^POS^ B cells. The identification of the increased expression of the IL2-STAT5 signaling pathway suggested a possible role for IL-15 since IL-2 and IL-15 share the same β (CD122) and γ (CD132) receptors in mediating signaling^47^ (**Figure 4C**). We tested this hypothesis by examining the relative abundance of both *IL15* and its receptor *IL15RA*. Although not identified as DEGs, both *IL15RA* (**Figure 4D**) and *IL15* (**Figure S.4**) transcripts were upregulated within RA-CCP^POS^ B cells relative to either RA-CCP^NEG^ or HA^POS^ B cells.

**Figure 4.**
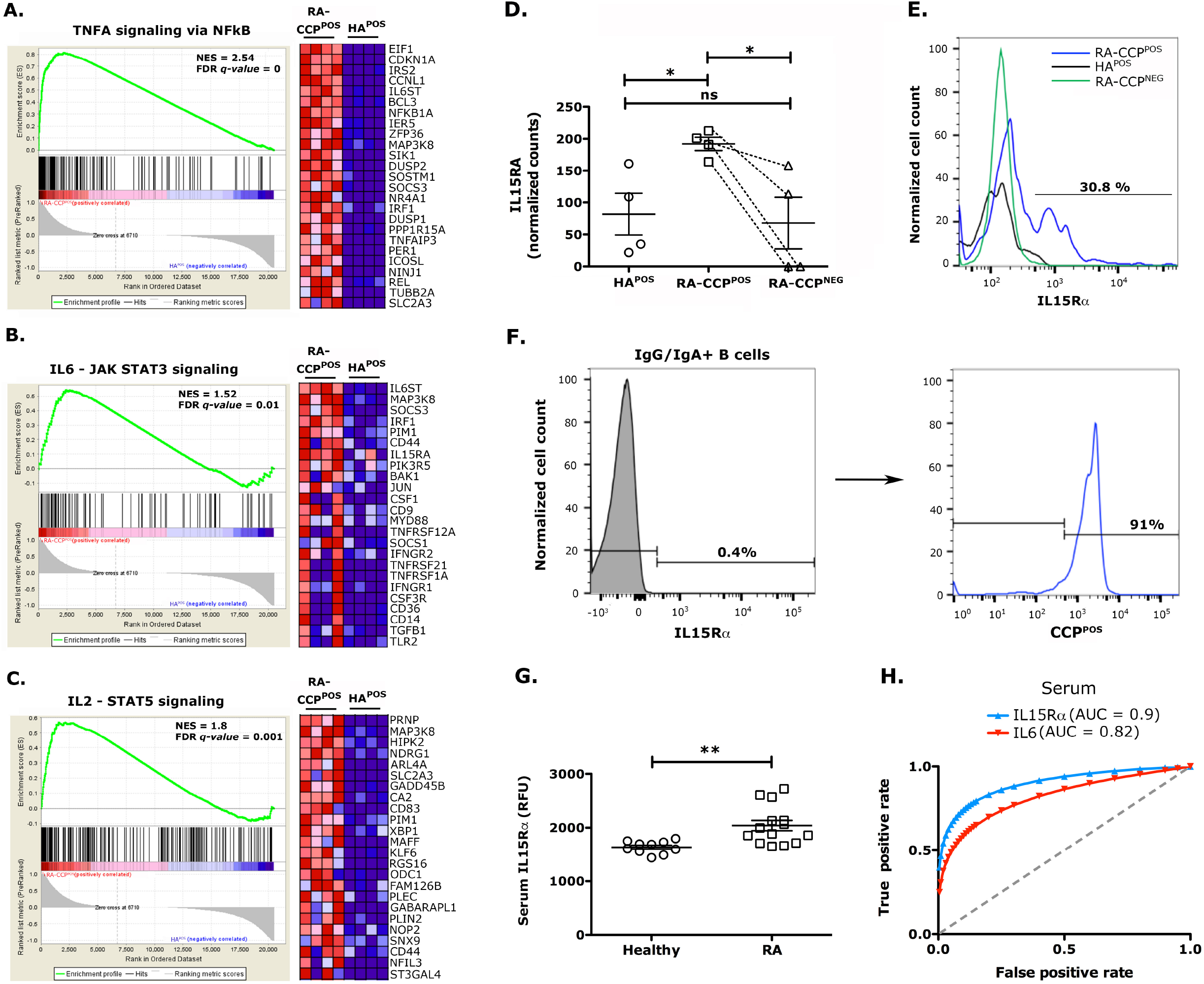
Proinflammatory TNF, IL15, and IL6 pathways are enriched in RA-CCP^POS^ B cells. (**A-C**) GSEA enrichment plots derived from pre-ranked DEGs of cytokine pathways of TNF, IL6, and IL2 indicating normalized transcript levels with a heat map of top twenty-five genes in the pathway for each sample. **D**. Normalized transcript’s count of gene *IL15RA* in RA-CCP^POS^, RA-CCP^NEG^, and HA^POS^ B cells. **E**. Representative flow-cytometric plot of surface expression of IL15Rα on RA-CCP^POS^, RA-CCP^NEG^, and HA^POS^ B cells (n = 4). **F, G**. Restricted expression of IL15Rα on RA-CCP^POS^ B cells when gated on IL15Rα positive total B cells or IgG/IgA B cells (n = 3). **H**. Serum concentration of IL15Rα protein in RA patients and healthy individuals evaluated by SOMAscan aptamer assay (RFU = relative fluorescence unit). **I**. Receiver operating characteristic curve (ROC) indicating the high predictive value of IL15Rα for RA with an area under the curve (AUC) value of 0.9 against IL6 (AUC = 0.82). * *p-value* < 0.05, ** *p-value* < 0.01 * *p-value* < 0.05. NES – Normalized enrichment score; FDR – False discovery rate. Statistical analysis in panels D and H is by Mann-Whitney U test.

In order to validate the elevated expression of IL15Rα, we utilized an independent cohort of RA patients (**Table S.3**) and performed *ex vivo* surface staining on B cells by flow cytometry. Consistent with the RNA-seq results, there was a distinct subpopulation of RA-CCP^POS^ B cells expressing IL15Rα in all of the patients tested while these were clearly absent in either the RA-CCP^NEG^ or HA^POS^ B cell populations (**Figure 4E**). Interestingly, within RA patients, >90% of IL15Rα expressing IgG/IgA B cells, were identified to be RA-CCP^POS^ B cells, thus indicating that IL15Rα expression is restricted and selective to this population (**Figure 4F**).

Since IL15Rα can also be converted to the soluble form by proteolytic cleavage by TNF-alpha converting enzyme (TACE)^48^, we next investigated the concentration of soluble IL15Rα (sIL15Rα) in RA patient’s cohorts. To systematically validate a large number of candidate proteins within cohorts of RA patient’s sera, including IL15Rα, we took advantage of the aptamer-based SOMAscan assay that can detect analytes at picomolar concentrations by utilizing very small volumes of biological samples (**Figure S.5**). The SOMAscan assay has been independently validated in a number of disease settings including musculoskeletal diseases^49^. In a cohort of RA patients (14 samples) compared to healthy donors (10 samples), we observed significantly higher concentrations of both IL-6 and IL-8 but not in a large panel of other soluble analytes including IL-1β or IL-17 (**Figure S.6**). When we evaluated sIL15Rα, we observed significantly higher concentrations in the sera of RA patients in comparison to healthy controls (**Figure 4G**). Lastly, we also looked at the predictive value of serum sIL15Rα determined by the SOMAscan assay and found that sIL15Rα displays both high sensitivity and specificity for RA patients in comparison to healthy controls (**Figure 4H**).

### EGFR pathways and molecular validation of AREG in RA B cells

AREG is a member of the epidermal growth factor family of ligands that signal through the EGFR receptors. It has been previously documented that TNF signaling in conjunction with IL-1β signaling can lead to the up regulation of AREG^48,50^. Our data supports a pivotal role for *AREG* in RA-CCP^POS^ B cells (**Figure 5A**), and the downstream target pathway, EGF-mediated signaling (**Figure 5B**) and the related NRG1 mediated signaling (**Table S.4**). Within the adaptive immune cell compartment, although the impact of AREG has been described on regulatory T cells, there are no known reports of B-cell specific expression. In order to validate our findings, we directly interrogated AREG expression within individual B cells on an independent cohort of RA patients by flow cytometry (**Table S.3**). As expected, we did not observe surface expression of AREG on RA-CCP^POS^ B cells *ex vivo* (data not shown) since AREG is secreted as soluble protein by TNF-activated B cells. In order to determine if AREG expression can be induced *in vitro* in B cells, we sought to mimic the nature of help afforded by T helper cells *in vivo* in RA. Accordingly, populations of RA-CCP^POS^, RA-CCP^NEG^, and HA^POS^ B cells were flow sorted and incubated with both soluble CD40L and IL-21 for 14 days. Under these conditions, both RA-CCP^POS^ and RA-CCP^NEG^ B cells showed induction of AREG in >80 *%* of cells (**Figure 5C**), and this was consistent with our RNA-seq data on these populations (**Figure 5A**). A tendency towards higher expression was observed when RA-CCP^POS^ and HA^POS^ B cells were compared for AREG expression but owing to the high variance in frequency of AREG expressing HA^POS^ B cell populations this change was not significant. Taken together, these findings suggest that under polarizing conditions, at least *in vitro*, B cells from RA patients act as a source of AREG, a molecule with a known ability to have a global impact on multiple cell types.

**Figure 5.**
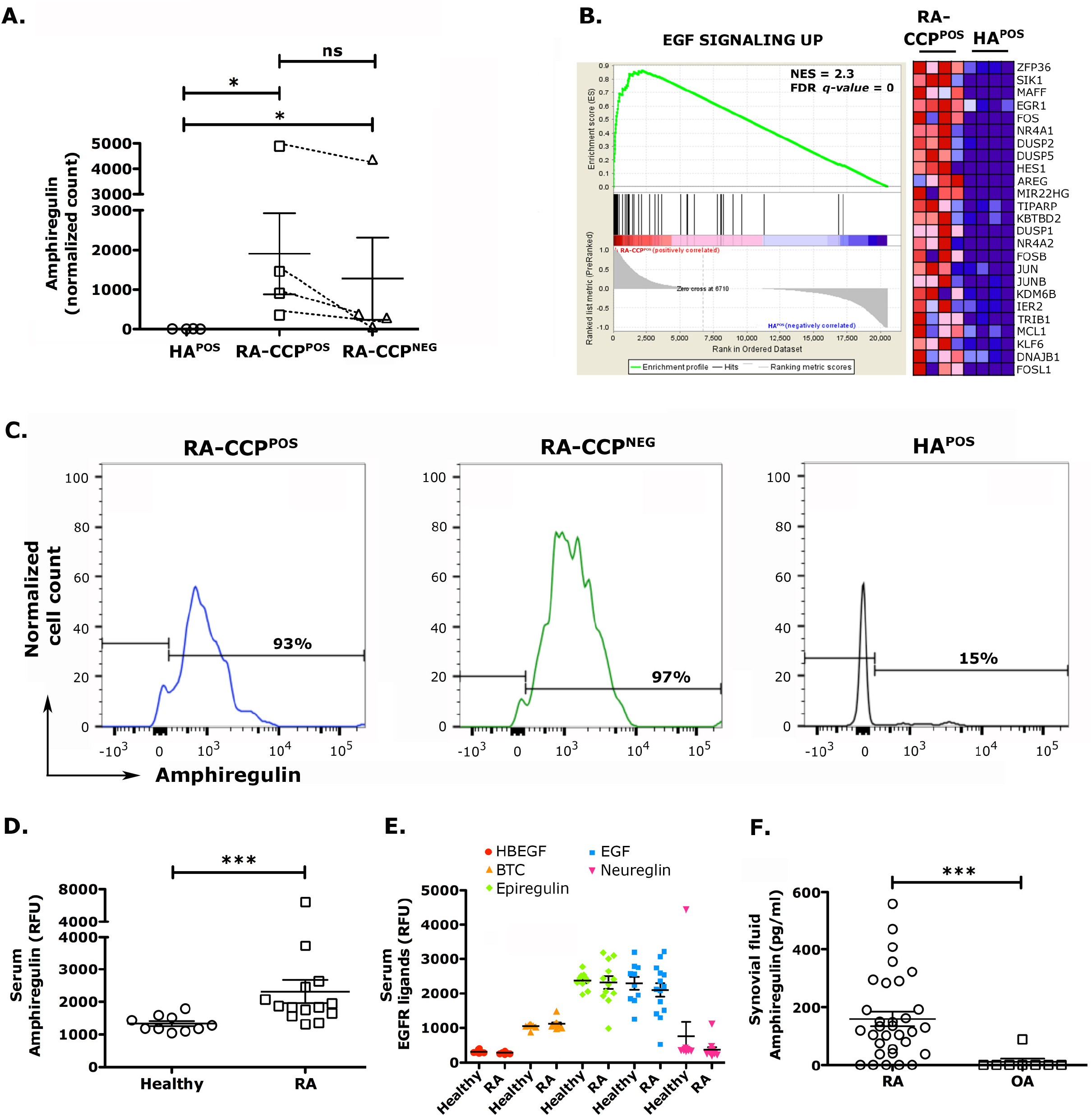
Amphiregulin and associated pathways are enriched in RA-CCP^POS^ B cells. **A.** Normalized transcript count of *AREG* in RA-CCP^POS^, RA-CCP^NEG^ and HA^POS^ B cells. **B**. GSEA enrichment plot of epidermal growth factor pathway with heat map indicating normalized transcript levels of top twenty-five genes in the pathway for each sample. **C**. Flow-cytometric evaluation of surface expression of amphiregulin on RA-CCP^POS^, RA-CCP^NEG^, and HA^POS^ B cells after *in vitro* expansion (n = 4). **D**. Serum concentration of AREG protein in RA patients and healthy individuals evaluated by SOMAscan aptamer assay. **E**. Serum concentrations of all the other EGF ligands identified by SOMAscan aptamer assay. **F**. ELISA on synovial fluid (SF) from patients with RA as compared to control osteoarthritis (OA). * *p-value* < 0.05, ** *p-value* < 0.01. Statistical analysis in panels A, D, E and F is by Mann-Whitney U test.

### AREG expression is elevated in serum and synovial fluid of RA patients

Among the different EGF ligands that are known to induce proliferation, growth, and differentiation by signaling through the EGFR receptors^51^, AREG plays a unique role in its ability to induce both cell proliferation and cellular differentiation upon receptor binding^52^. We evaluated the concentrations of AREG and the other EGF ligands in the sera of RA patients with the SOMAscan array, as outlined above. In this cohort of samples, we observed a significantly higher concentration of AREG (**Figure 5D**) but none of the other EGF ligands like heparin-binding EGF-like growth factor (HBEGF), betacellulin (BTC), epiregulin, EGF and neuregulin (**Figure 5E**). We also observed a high sensitivity and specificity of serum AREG in predicting the occurrence of RA (**Figure S.7**). Since RA is a disease with localized inflammation, we further evaluated the abundance of AREG in the synovial fluid (SF) from RA patients (32 samples) and compared it with the SF from the non-inflammatory type osteoarthritis (OA, 8 samples). Our ELISA results showed that the levels of AREG were significantly higher in RA as compared to osteoarthritis (OA) patients, wherein only one patient’s SF tested positive for AREG (**Figure 5F**).

### Amphiregulin promotes migration and proliferation of fibroblast-like synoviocytes (FLS) *in vitro*

One of the hallmarks of RA pathogenesis is the activation of the FLS, which is characterized by a tumor-like aggressive phenotype within the joints^53^. We tested the ability of AREG to increase the invasiveness of human RA-FLS and observed that AREG promoted the increased migration of FLS in wound healing assays, in comparison to IL15 or controls (**Figure 6A, B**). We confirmed the expression of EGFR on RA-FLS, suggesting that the increased invasiveness is likely due to increased AREG-EGFR signaling (**Figure 6C**). By contrast, however, a lack of RA-FLS migration in the presence of IL15 could be attributed to the low surface expression of its cognate receptor IL15Rα (data not shown). We also observed that RA-FLS proliferate more in the presence of AREG in comparison to controls (**Figure 6D**) or IL15 (data not shown). Taken together, our results thus indicate that high-levels of AREG in RA patient’s blood and SF could be responsible for an aggressive phenotype of FLSs in these patients.

**Figure 6.**
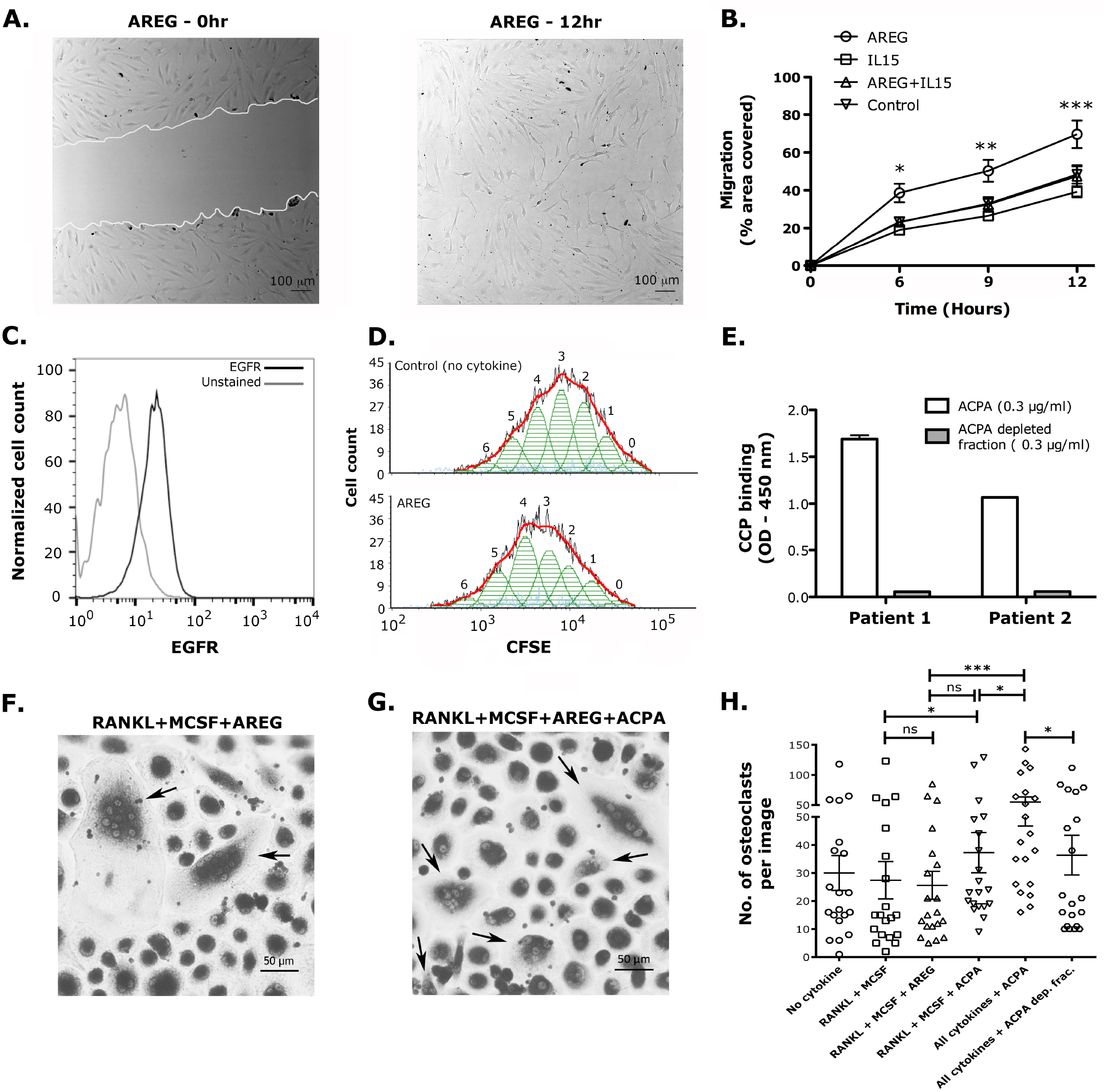
Amphiregulin enhances cell migration and proliferation of fibroblastlike synoviocytes and synergizes with ACPA in mediating osteoclast differentiation. **A**. Scratch assay depicting cell migration in presence of AREG (white line indicates the boundary of the scratch). **B**. Migration of FLS monitored at different time points under different culture conditions represented as the scratch area covered. **C**. Expression of EGFR on FLS; gray line represents unstained cells. **D**. The proliferation of FLS in the presence or absence of AREG visualized as dilution of the CFSE staining; numbers indicate different generations of AREG-treated FLS in comparison to untreated cells; experiments performed in triplicates. **E**. ELISA to evaluate the purity of ACPA enrichment from the plasma of two RA patients against CCP. **F**. Differentiation of osteoclasts from blood monocytes in the presence of MCSF, RANKL, and AREG. **G**. Osteoclasts differentiated in the presence of all cytokines plus ACPA; arrows indicate differentiated osteoclasts with ≥ 3 nuclei. **H**. Number of osteoclasts (TRAP^+^ cells with ≥ 3 nuclei) per image under different culture conditions; a total of ten images from random field of views per well were taken at 10x magnification in duplicate culture conditions. * *p-value* < 0.05, *** *p-value* < 0.001. Statistical analysis in panel B is by two-way ANOVA and in panel G by Mann-Whitney U test. * represents significance between AREG and control.

### Functional synergy between AREG and ACPA in mediating osteoclastogenesis

The role of AREG in promoting osteolytic activity has been detailed as one of the mediators of bone metastases in breast cancers^54^. Independently, the role of ACPA in enabling osteoclast differentiation has been described in RA^5^. We thus interrogated the functional synergy between these molecules in mediating osteoclastogenesis. First, we purified the ACPA antibodies by binding to a CCP-functionalized column (**Figure 6E**). Next, we evaluated the ability of each of these molecules either by themselves or in combination to promote the differentiation of multi-nucleated Tartrate-resistant acid phosphatase (TRAP) positive osteoclasts from monocyte precursors. Although AREG by itself did not significantly increase the differentiation of osteoclasts; the number of osteoclasts was observed to be higher in co-cultures with AREG and ACPA in comparison to all other conditions tested (**Figure 6F-G**). Our findings thus suggest that AREG can synergize with ACPA in the differentiation of osteoclasts from blood monocytes.

### cAMP in RA-CCP^POS^ B cells

The secondary messenger molecule cyclic-AMP (cAMP) is a major metabolic regulator of immune cell fate and effector functionality^55^. In comparison to HA^POS^ B cells, within the RA-CCP^POS^ B cells, we observed an enrichment of a number of transcription factors (TFs) or activators that globally activate or repress gene transcription by recognizing the cAMP response element (CRE)^56^. These included the activating transcription factors (*ATF4* and *ATF5*)^57^, AP-1 transcription factors (*Fos, JunB*, and *JunD*)^58^, Cbp/P300 interacting transactivator with Glu/Asp-rich carboxy-terminal domain 4 (*CITED4*)^59^, and Period circadian protein homolog 1 (*PER1*)^60^ (**Figure 2E**). The stability and half-life of intracellular cAMP are modulated by the phosphodiesterase (PDE) family of enzymes^61^. Consistent with the expression of cAMP within these RA-CCP^POS^ B cells, *PDE4B* which is the major PDE found in immune cells was identified as a DEG with increased expression within this cellular population^62^. GSEA analysis of the TF targets (TFT) confirmed the enrichment of both CREB and ATF targets (**Figure S.8**). The differential expression of the cAMP pathways was also observed in the RA-CCP^POS^ B cells in comparison to the RA-CCP^NEG^ B cells (**Figure 3A**).

## Discussion

The role of RA-CCP^POS^ B cells as a source of autoantibodies contributing to disease pathology in RA has been extensively studied but the functional programs that help define the biology of the cell have remained elusive. One of the challenges we encountered was the robust and reliable identification of RA-CCP^POS^ B cells. Although citrullinated antigen-specific cells in the synovial fluid have been documented to comprise up to 25% of the total B-cell population in the joints^63^, CCP-specific B cells in blood account for <0.1% of total peripheral B cells of RA patients^19^. We successfully designed a sensitive flow sorting method using peptide-streptavidin tetramers, an approach similar to another recently published report but with one major difference^19^, we sorted both the IgG^pos^ and IgA^pos^ CCP^POS^ B cells, owing to the fact that the antibody response against citrullinated proteins in RA comprises both of IgG and IgA type immunoglobulins^64^.

Our study outlines specific transcriptional and functional differences in RA-CCP^POS^ B cells within RA patients in comparison to either RA-CCP^NEG^ B cells from the same patients or HA^POS^ B cells from healthy donors. Perhaps the least surprising finding is the well-known cytokine-mediated inflammatory pathways involving TNF and IL-6. There are multiple inhibitors and biologics targeting these molecules, their receptors or downstream signaling molecules that are already approved or in various stages of approval (**Table 2**). Comparisons of the RA-CCP^POS^ and RA-CCP^NEG^ B cells suggests that the impact of these cytokines were likely global, influencing all B cells within these patients.

**Table 2.**
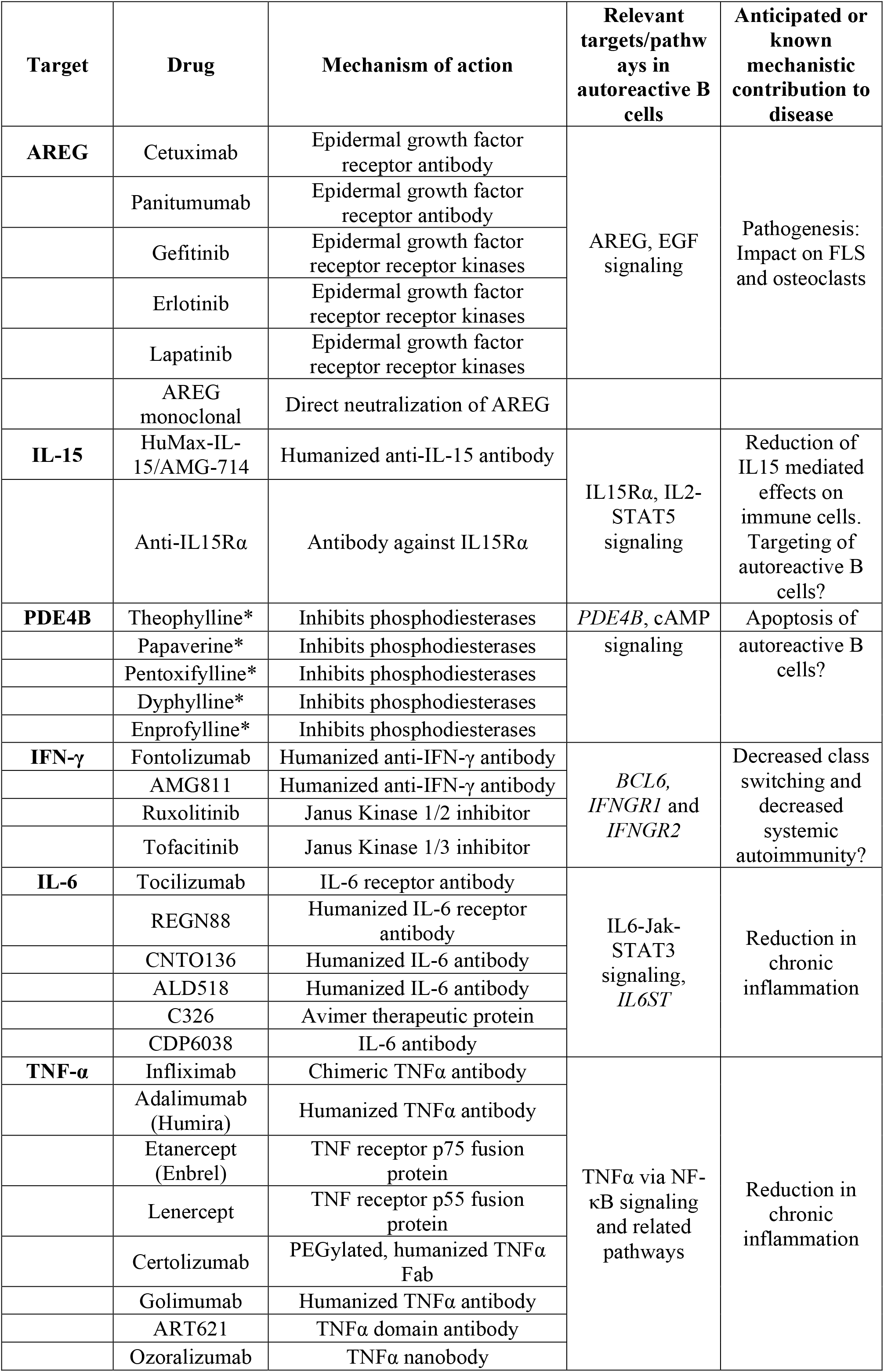
Drug targets and pathways identified based on RNA-seq on autoreactive B cells in RA.

Based on the RNA-seq data, we validated that the expression of the IL15Rα protein on the cell surface was restricted to the RA-CCP^POS^ B cell population, and confirmed that increased levels of sIL15Rα can be detected within the blood of an independent cohort of RA patients. IL-15 is a four-helix bundle glycoprotein and forms a complex with its private receptor, IL-15Rα, and this complex can then bind to the shared IL-2 β-receptor (IL-15Rβ, CD122) and the common cytokine receptor γ chain (CD132)^65^. IL-15 derived signaling is known to promote the proliferation of Th1/Th17 cells, and the proliferation and immunoglobulin (Ig) production from B cells^66,67^. With regards to RA, high concentrations of IL-15 in circulation and IL-15Rα in the synovium are known to be associated with increased disease activity^68^. Genome-wide association studies (GWAS) have identified single-nucleotide polymorphisms (SNPs) in the IL-15 gene that are correlated with bone erosion and poor functional outcomes in RA^69^. IL-15 has been targeted in phase I-II clinical trials by using HuMax-IL15 (**Table 2**), a human monoclonal antibody that inhibits the bioactivity of IL-15, and showed modest improvements in disease activity^70^. Synthetic DMARDs including tofacitinib (Janus kinase inhibitor) inhibit pathways that are also downstream of IL-15Rα derived signaling^65^. Our data suggests that the targeting of IL15Rα might provide therapeutic benefit in RA either by directly targeting the RA-CCP^POS^ B cells or by the elimination of sIL15Rα in the blood of RA patients. In this regard, it is worth emphasizing that sIL15Rα is known to increase the potency of IL15 function by 100 fold when complexed together, and enables signaling in cells that might otherwise lack IL15Rα^71^. One of the novel observation of our study is the abundance of cAMP-mediated pathways within the RA-CCP^POS^ B cell populations. cAMP is a vital secondary messenger that plays an essential role in cellular response to many hormones, GPCRs and growth factor receptors^72^. The intracellular concentrations of cAMP are determined by the activity of two opposing enzyme families, the adenylyl cyclases (AC) and cyclic nucleotide phosphodiesterases (PDEs). The best-characterized effector of cAMP is protein kinase A (PKA) that phosphorylates a large number of cytosolic and nuclear proteins^73^. Transcripts of PKA are directly regulated by the phosphorylation of the transcription factors cAMP-response element-binding protein (CREB), a cAMP-responsive modulator (CREM), and ATF1. It is well-documented that CREB is activated downstream of BCR signaling, and plays important roles in B cell survival and function^74^. The increased signaling through the BCR of autoreactive B cells might provide one explanation for the increase in cAMP signaling pathways documented here. The synthetic DMARD methotrexate inhibits the enzyme dihydrofolate reductase and related enzymes, ultimately leading to the release of adenosine into the extracellular environment^75^. This, in turn, leads to increased cAMP levels by acting through the adenosine receptors, and since all our RA-CCP^POS^ B cells were isolated from RA patients that had undergone methotrexate therapy, it is plausible that the increased cAMP signaling might be linked to therapy. Despite this, the specific increase of cAMP within the RA-CCP^POS^ B cells, and the identification of *PDE4* as a DEG suggests that inhibition of PDE4 could be explored as a potential mechanism to induce apoptosis within RA-CCP^POS^ B cells.

Our data for the first time also revealed high expression of EGFR ligand AREG on autoreactive B cells, which was also the most differentially expressed gene in RA-RA-CCP^POS^ B cells as compared to HA^POS^ B cells. We confirmed the surface expression of AREG on B cells under inflammatory conditions, identified AREG as a candidate biomarker in RA and have evaluated the functional impact of AREG on the cellular effectors of RA. AREG is expressed as a membrane-bound molecule, which is activated upon proteolytic cleavage by TACE^48^, thereby acting in an autocrine or paracrine manner. In addition, pro-AREG can also activate cells expressing EGFR in a juxtacrine manner^76^. Binding to the EGFR receptor leads to homodimerization or heterodimerization of the EGFR receptor with ErbB2, ErbB3, and ErbB4, resulting in the activation of a signaling cascade that influences the cell’s survival, proliferation and motility^76^.

Overexpression of AREG is best characterized in the context of tumors where it has been shown to promote the migration and invasiveness of cancer cells^54^. More recently, immunohistochemistry has confirmed the expression of AREG within the synovial tissue of RA patients^48^, and AREG has also been associated with the pathogenesis of other autoimmune diseases like psoriasis^77^. Indeed, mouse models of psoriasis display synovitis and treatment with mAbs targeting AREG alleviates inflammation^78^.

The induction of AREG expression is governed by the cytokines TGFβ, TNFα, and IL1^β48,50^, known to be abundant in RA synovial fluid and plasma^79^. As our results outline, the activation of the TNFα and TGFβ pathways (**Figure 4A, S.9**) may allow RA-CCP^POS^ B cells to secrete AREG along with other pro-inflammatory cytokines. Furthermore, as our data illustrates, AREG can directly promote the proliferation and invasiveness of FLS. As outlined by elegant studies in breast cancer metastasis models, AREG is also known to increase the differentiation and activity of osteoclasts from PBMCs in the presence of RANKL and M-CSF^54^, both of which are abundantly available in the synovial compartment of RA patients^80,81^. Our *in vitro* data further advances the ability of AREG to act in concert with ACPA to enhance the differentiation of osteoclasts from blood monocytes. We thus propose an integrated model for the many different impacts of RA-CCP^POS^ B cells in RA based on both our data and on the multi-functional roles of AREG in other disease models (**Figure 7**). Through this study, we are reporting for the first time a comprehensive transcriptome profile of RA-CCP^POS^ B cells in RA. To the best of our knowledge, this is also the first report on whole transcriptome profiling of antigen-specific B cells in any autoimmune disorder. Our results portray B cells as not merely autoantibody producers but as a source of diverse molecules which can influence proliferation, differentiation, and activation of other pathogenic cells types. We anticipate that these data will serve as a foundational data set for investigating multiple hypotheses on the roles of B cells in RA and other autoimmune disorders, and enable drug discovery and validation based on the biology of RA-CCP^POS^ B cells in RA.

**Figure 7.**
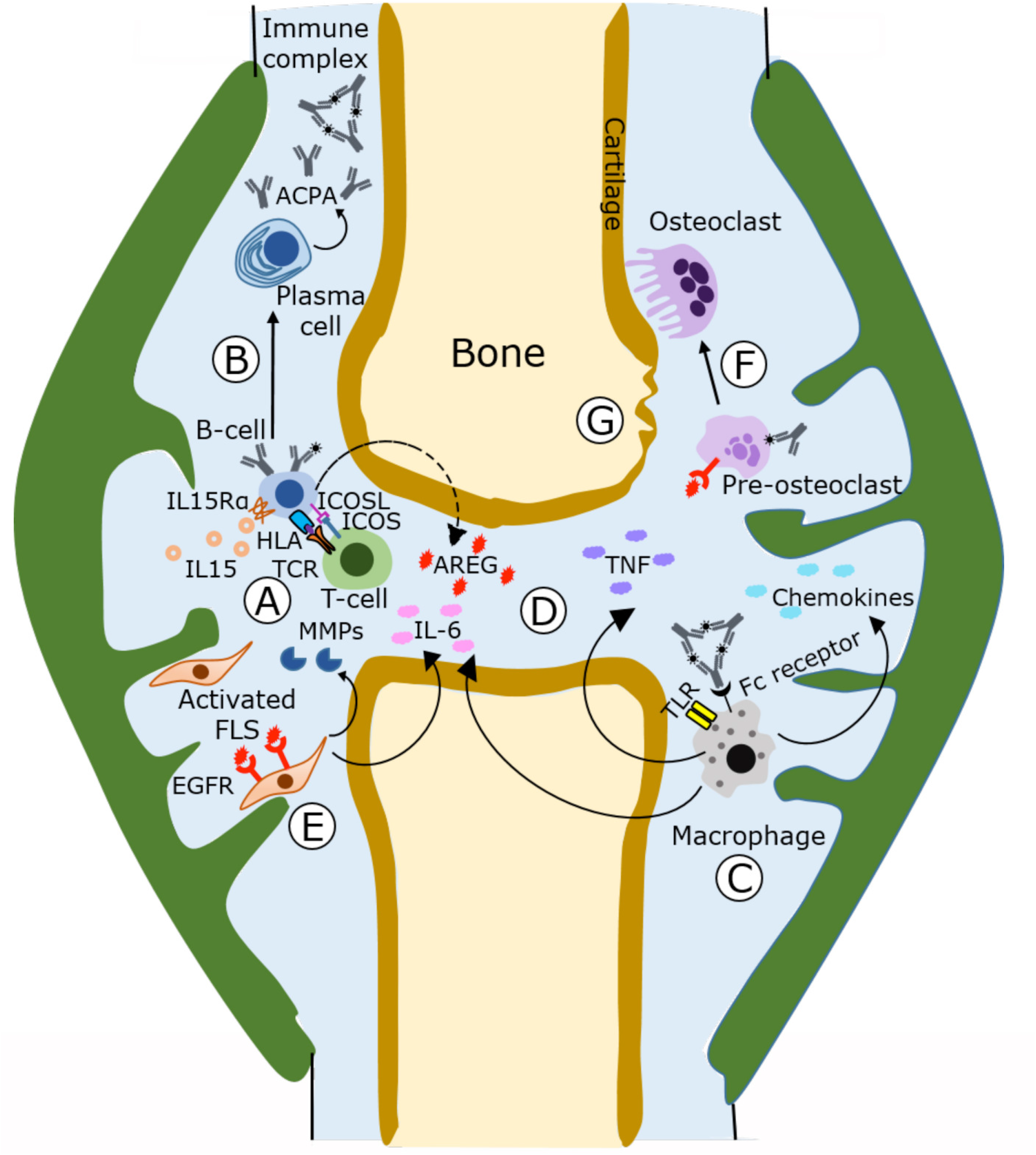
A proposed integrated model for the multi-faceted roles of B cells in RA. (**A-B**) Activated autoreactive B cells likely with the help of T cells, in the presence of IL15, enable the somatic hypermutation and differentiation of B cells into plasma cells which in turn secrete ACPA and other autoreactive antibodies. (**C**) ACPA in the presence of citrullinated antigen form immune complexes that can activate macrophages through the Fc receptors, and in concert with TLR stimulation, lead to the secretion of chemokines, TNF and IL-6. (**D**) This establishes a systemic inflammatory loop which directly impacts B cells and can enable the secretion of AREG. Additionally, it is likely that other immune and non-immune cells also secrete AREG under these conditions. (**E**) AREG (signaling through EGFR) along with the inflammatory microenvironment enable the proliferation and activation of FLS which in turn secrete MMPs to increase invasive destruction. (**F-G**) ACPA along with AREG can promote the increased differentiation of pre-osteoclast into osteoclasts and mediate bone destruction. Since all these cascades are cyclic in nature, more mechanistic studies will have to be undertaken to identify the early events that seed the cascade in contrast to the molecules and events that serve to amplify it.

We also recognize that while our report represents an important first step, further studies are required to gain a much deeper understanding of autoreactive B cells in autoimmune biology. Since our studies profiled CCP reactive B cells *ex vivo* from treated patients, it is not clear how treatment can impact the profiles described here. For this reason, it would be ideal to study new onset treatment naïve patients and perform RNA-seq on B cell isolated from these patients. At the other end of the spectrum, since RA is an inflammatory disorder, and since the synovial compartment is a primary region of disease activity and pathology, it will be interesting to profile the autoreactive B cells isolated directly from the synovial mononuclear cells of advanced RA patients. Nonetheless, with the aid of the *ex vivo* sorting method that we have utilized, these questions can be addressed with careful selection of the appropriate patient cohorts. Similarly, the relationships between IL15, sIL15Ra, and AREG within the synovial compartment, and how they influence the expression and secretion of each other and other signaling cascades, needs better definition.

## Acknowledgements

This publication was supported by the NIH (R01CA174385), CPRIT (RP130570); MRA Stewart-Rahr Young Investigator Award to NV; Welch Foundation (E1774) and the UAB RADAR fund. We would like to acknowledge the UH Seq-n-edit core for RNA-seq, Intel for the loan of computing cluster, and the UH Center for Advanced Computing and Data Systems (CACDS) for high-performance computing facilities.

## Author contributions

Designed the research: AM, NV, CM

Performed the experiments – AM, SA, SS, GR, AA

Analyzed the data – AM, XY, SA, DP, JTA, JR, CC, PQ

Contributed research material – KW, SLB, AA, SKA, CM

Wrote the paper – AM, NV, CM

## Material and methods

### Patients

Blood (15 ml) was aspirated in heparin vacutainer tubes (BD Biosciences) from RA patients after informed consent under IRB approved protocols at BCM, UAB and UH. Patient information is summarized in Table 1. All RA patients met 1987 ARA (now ACR) classification criteria for RA^82^.

### Flow-sorting of CCP-specific B cells from patients’ blood

Peripheral blood mononuclear cells (PBMC) from RA patients were stained for B-cell markers with fluorochrome-labeled antibodies, anti-CD19, IgM, and IgD together with T-cell marker using anti-CD3 antibody (Biolegend) according to manufacturer’s instructions. Biotinylated-cyclic citrulline peptide I (Anaspec) was then added (1 μg/million PBMCs) followed by labeling with fluorochrome-labeled streptavidin (1 μg/ml). Subsequently, cyclic arginine peptide (Anaspec) was also added (1 μg/million PBMCs) and labeled with fluorochrome-labeled streptavidin (1 μg/ml), to exclude B-cell crossreactive to both antigens. All staining steps were performed on ice for 30 min each and cells were washed/suspended in cold PBS-FBS 2%. Further, CD19^pos^IgM/IgD^neg^ B cells (IgG/IgA^pos^) were gated and the three cell populations CCP^POS^AP^NEG^, CCP^NEG^AP^POS^, and CCP^NEG^AP^NEG^ were sorted for validation of cell purity. A total of 125-360 RA-CCP^POS^ B cells were obtained of 10-15 million PBMCs from each patient. The sorted cells were then grown in 96-well tissue culture plates with 1x10^5^/well of 3T3 fibroblast cells secreting msCD40L (NIH AIDS reagent program), 100 ng/ml of IL-21 (Peprotech) and 5 μg/ml of anti-APO1 antibody (eBiosciences) for a period of 14 days with half media change once after 7 days. Thereafter, cell purity was assessed by validating the specificity of IgG from the culture supernatants against CCP.

### Supernatant CCP-ELISA

The 96 well maxisorp plates (Nunc) were coated with streptavidin (2 μg/ml), followed by blocking with PBS-BSA 1% and then incubated with/without the peptides (4 μg/ml). Supernatants from each cell population were mixed 1:1 in blocking buffer (PBS-BSA1%) and then incubated in triplicates in wells coated with: (a) streptavidin only, (b) streptavidin incubated with AP, and (c) streptavidin incubated with CCP. After subsequent washing, the presence of specific binding of immunoglobulins with the antigen-peptides was detected by adding HRP-labeled anti-human IgG antibody (Jackson) at a dilution of 1:10000, followed by colorimetric estimation at OD – 450 nm after the addition of TMB substrate and stopping the reaction by 2N H_2_SO_4_.

### ELISA of HA plasma

Plasma antibody titers of donors against HA from Influenza A H3N2 (A/Texas/50/2012) strain (Sinobiologicals) used to vaccinate the individuals, was measured by coating microtiter plates with HA (1 μg/ml), followed by blocking with PBS-BSA 1%. Thereafter, donor plasma was added (1:1000 dilution) and antibody titers were detected by using anti-human IgG-HRP antibodies (1: 10,000 dilution) followed by addition of TMB substrate and stopping the reaction by 2N H_2_SO_4_.

### Isolation of HA-specific B cells

The HA-specific B cells were sorted using recombinant HA from the same Influenza A strain as listed above. B-cell staining was performed with the same panel of antibodies as used for the RA-CCP^POS^ B cells, followed by incubation with HA (1 μg/million PBMCs). The cells were then incubated with rabbit anti-HA antibody (Sinobiologicals) at a concentration of 2 μg/ml, followed by incubation with 2 μg/ml of donkey anti-rabbit antibody (Biolegend). All incubations were performed for 30 minutes on ice. Similar to RA-CCP^POS^ B cells, the purity of flow-sorted cells was assessed by culturing HA^POS^ and HA^NEG^ cells from each individual with 3T3 fibroblast cells, IL-21, and anti-APO1 antibody. The specificity of sorted B cells was assessed by performing ELISA on supernatants obtained after 14 days of culture essentially as described above.

### Preparation of cDNA library and RNA sequencing

Total RNA was obtained by sorting antigen-specific B cells (350-1,000 total cells) *ex vivo* directly into the 100 μl of cell lysis buffer provided in the RNA isolation kit (Macherey-Nagel, RNA-XS). Further, cDNA libraries were synthesized using the commercially available SMART-Seq Ultra Low Input RNA kit (Clontech), as per the manufacturer’s protocols. After preparation of cDNA libraries, they were first tagmented and then barcoded by indexing primers using the Nextra XT kit (Illumina). The libraries were pooled and a 76bp paired-end sequencing was performed on an Illumina HiSeq3000 sequencer to yield a minimum of 17.4 million reads per library (range = 17.4 – 37.3 million).

### Bioinformatic pathway and gene discovery

Initial quality control of the sequencing data was performed by using Cutadapt to remove the adapter sequences from the 3’ end, followed by trimming of the first 12 bases of all reads at 5’ end to address the nonuniform distribution caused either by biased selection of reads or contamination of other sequences. Reads shorter than 20bp were then dropped to avoid potential mapping to multiple locations on the reference genome. After performing the quality control steps, transcripts were aligned to the human reference genome UCSC hg19 using TopHat2. The mapped genes were annotated using R package “GenomicAlignments” and gene annotation information in R package “TxDb.Hsapiens.UCSC.hg19.knownGene”. Thereafter, differentially expressed genes (DEGs) were identified using DEseq2 tool that tests for differential gene expression based on a model using the negative binomial distribution. The differentially expressed pathways were identified by the “GAGE” package using the gene sets downloaded from the “Molecular Signature Database” of the Broad Institute and also by using the Gene Set Enrichment Analysis (GSEA) software.

### Measurement of AREG in RA patient’s synovial fluid (SF)

The levels of AREG in patient’s SF were measured using the commercially available kit (R&D systems). A total of thirty-two RA and eight osteoarthritis (OA) patient’s SF was diluted 1:1 in assay diluent and was subjected to ELISA for the detection of AREG.

### Somamer assays

Serum was assayed using the standard protocol for the SOMAscan assay (Somalogic Inc.) targeting 1,310 specific human proteins. All samples were diluted to 40%, 1%, and 0.005% and incubated with SOMAmers (Slow Off-rate Modified Aptamers) for 3.5 hours at 28°C. The SOMAmers are conjugated to beads by a photocleavable linker and to a fluorophore that is used for final detection. After a series of washes with an assay buffer of physiological pH and salt to remove nonspecific binding, the remaining bound proteins were tagged using an NHS-biotin reagent. The aptamer-protein complexes were photocleaved from the beads by UV at a peak wavelength of 365nm and attached to streptavidin-coated magnetic beads. Using a high pH denaturing wash with a chaotropic buffer containing 12 SOMAmer hybridization controls, the SOMAmers bound to their cognate proteins were eluted and quantified using a custom DNA microarray and SureScan Microarray Scanner (Agilent Technologies). Raw data was sent to Somalogic to remove within-run hybridization variation and perform median normalization across samples to remove assay biases. The 40% dilutions of two RA samples were removed from analysis due to precipitates in the serum and all samples passed Somalogic’s quality control guidelines. R’s stats package was used for group statistics and normalization.

### Evaluating the surface expression of AREG and IL15Rα on B cells by flow cytometry

To identify the surface expression of AREG *ex vivo*, the cells were initially stained with 2μg/ml of primary mouse anti-AREG antibody (Sigma-Aldrich) followed by fluorochrome-labeled secondary anti-mouse IgG and fluorochrome-labeled antibodies against human CD19 and IL15Rα (Biolegend) according to manufacturer’s instructions.

### ACPA purification

Patient’s serum was diluted 1:10 in PBS and incubated overnight at 4°C with protein-G coated agarose beads. After incubation, the flow-through was removed and the column was washed thrice with PBS-Tween 0.1 %. The IgGs were then eluted using 0.1 M glycine (pH – 2.7) and neutralized with 1M Tris solution followed by buffer exchange with PBS. To isolate ACPA from total IgG, biotinylated CCP (0.5 mg) was incubated with 250 μl of agarose beads coated with streptavidin (Thermo Fisher) for 1hr at RT and then the unbound CCP peptide was washed with PBST-0.1%. Further, total IgG was first incubated with streptavidin (10 μg/ml) for 1hr, prior to incubation on the CCP-coated beads, to prevent isolation of streptavidin specific IgGs. ACPA was then isolated from total IgGs after overnight incubation at 4°C using the same elution strategy as described above.

### Osteoclast differentiation

Peripheral blood mononuclear cells were isolated from a healthy donor and 400,000 cells were plated per well on a poly-lysine coated glass bottom 24-well plate (MatTek Corporation). After six hours, non-adherent cells were washed using PBS and the cells were allowed to differentiate in serum-free X-VIVO 10 media (Lonza) with 1 ng/ml RANKL (Biolegend) and 10 ng/ml MCSF (R&D Systems) under different combinations of culture conditions with 100 ng/ml AREG (R&D Systems), ACPA (50 ng/ml) and ACPA-depleted fraction (50 ng/ml) for 14 days, with periodic change of media every 3rd day. Osteoclasts were labeled using a commercial kit (Sigma-Aldrich) for TRAP positivity and nuclear staining and visualized under an inverted microscope at 20X magnification. Images were taken from ten different fields of views per well from each of the duplicate culture conditions. Osteoclasts were counted as TRAP+ cells with ≥ 3 nuclei.

### Fibroblast-like synoviocyte migration assay

Human RA-FLS patient (CELL Applications, INC.) were grown to 100% confluency in 12-well plates in synoviocyte growth medium (CELL Applications, INC.). A scratch was created using a 200μl tip and the unbound cells were removed by washing with PBS. Cells were then incubated in synoviocyte basal media (CELL Applications, INC.) supplemented with 2% FBS under four different culture conditions, 100 ng/ml of AREG, 100 ng/ml of IL15 (Biolegend), AREG plus IL15 and media alone. Each culture condition was performed six times and cell migration was visualized using an inverted microscope (Carl Zeiss AG) at 10x (0.3 NA) magnification. Images were taken at time points 0hr, 6hr, 9hr, and 12hr. Migration of FLS was measured using the MRI wound healing tool in Image J.

---

**S. Figure 1.**
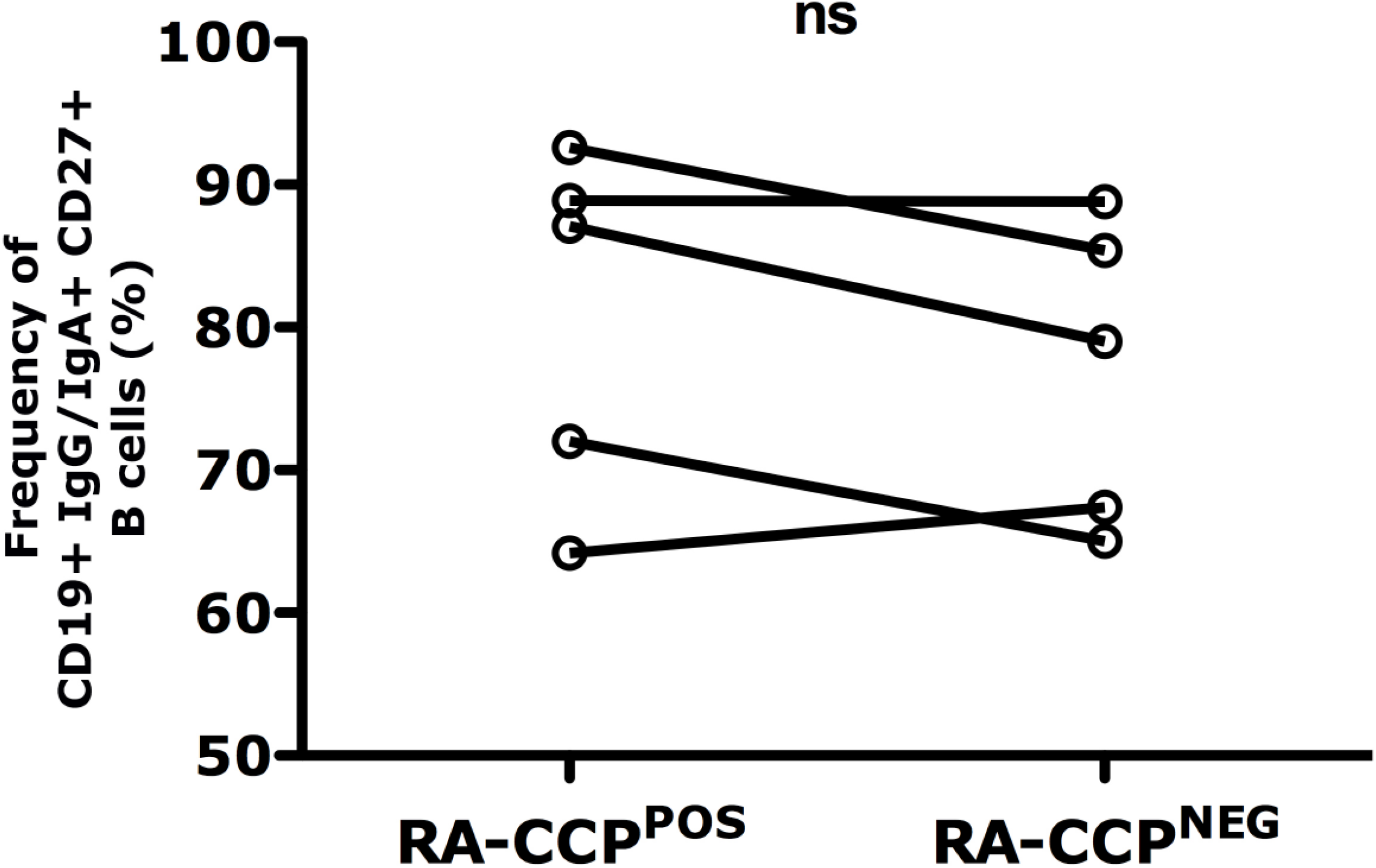
RA-CCP^POS^ and RA-CCP^NEG^ B cells are primarily of the memory phenotype. B cells from RA patient’s blood gated sequentially as CD19^+^ IgG/IgA^+^ CCP^POS^/CCP^NEG^ and memory phenotype defined as CD27^+^. Data from five paired RA samples includes four RA patients from whom RNA sequencing was performed. Statistical analysis is by paired t-test.

**S. Figure 2.**
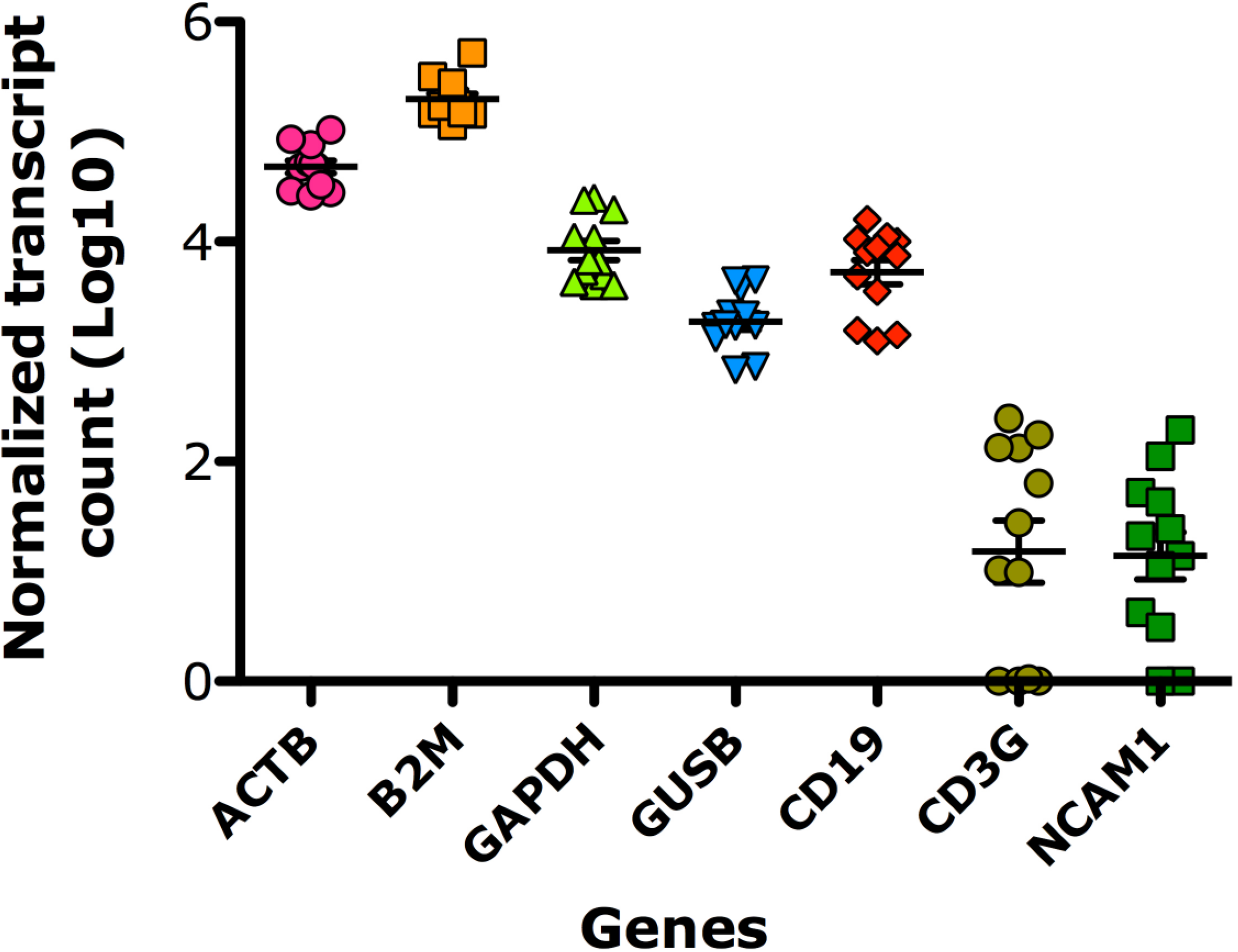
Quality control of RNA sequencing. Normalized transcript counts of house-keeping genes and immune-cell specific markers.

**S. Figure 3.**
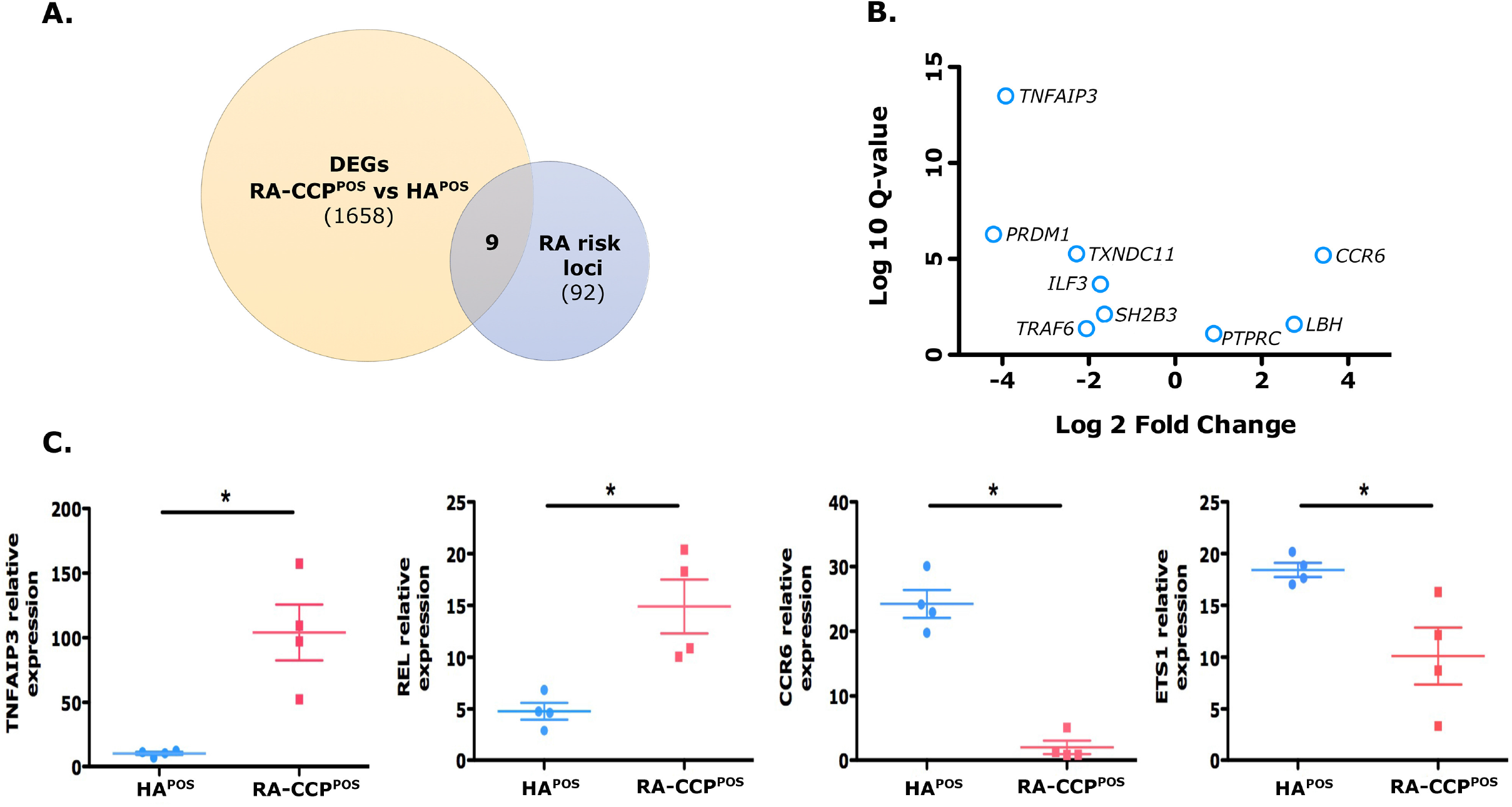
DEGs from RA-CCP^POS^ B cells overlap with few known risk loci of RA. **A.** Venn diagram of genes overlapping between DEGs and known risk loci of RA. **B.** Volcano plot displaying log-transformed *Q-value* and fold change expression of genes documented as risk loci in RA-CCP^POS^ and HA-specific B cells, **C**. Comparative normalized gene expression of risk loci genes in RA-CCP^POS^ and HA-specific B cells of each donor. * *p-value* < 0.05. Statistical analysis in panel C is by Mann-Whitney U test.

**S. Figure 4.**
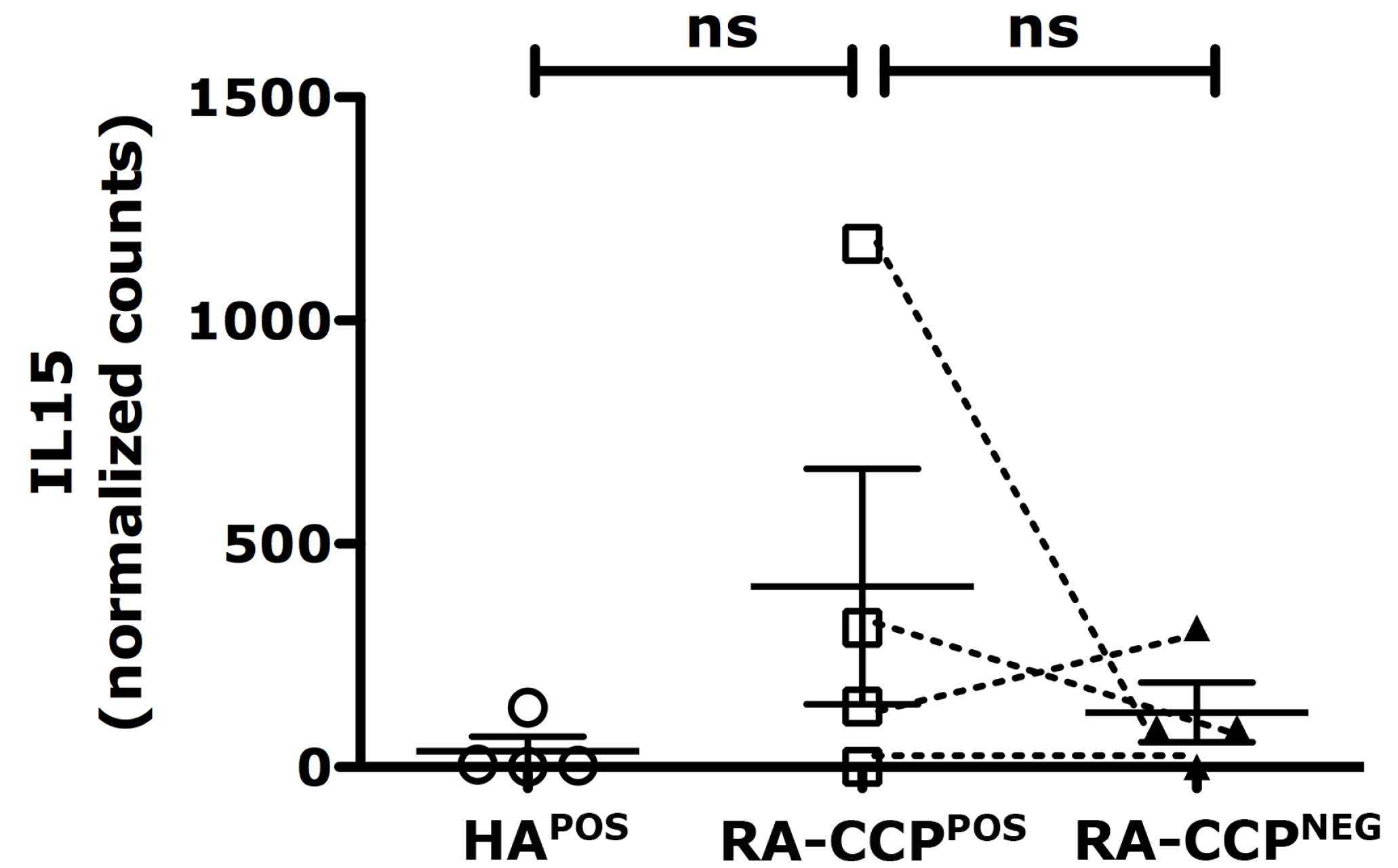
Normalized transcript counts of *IL15* in RA-CCP^POS^, RA-CCP^NEG^, and HA^POS^ B cells.

**S. Figure 5.**
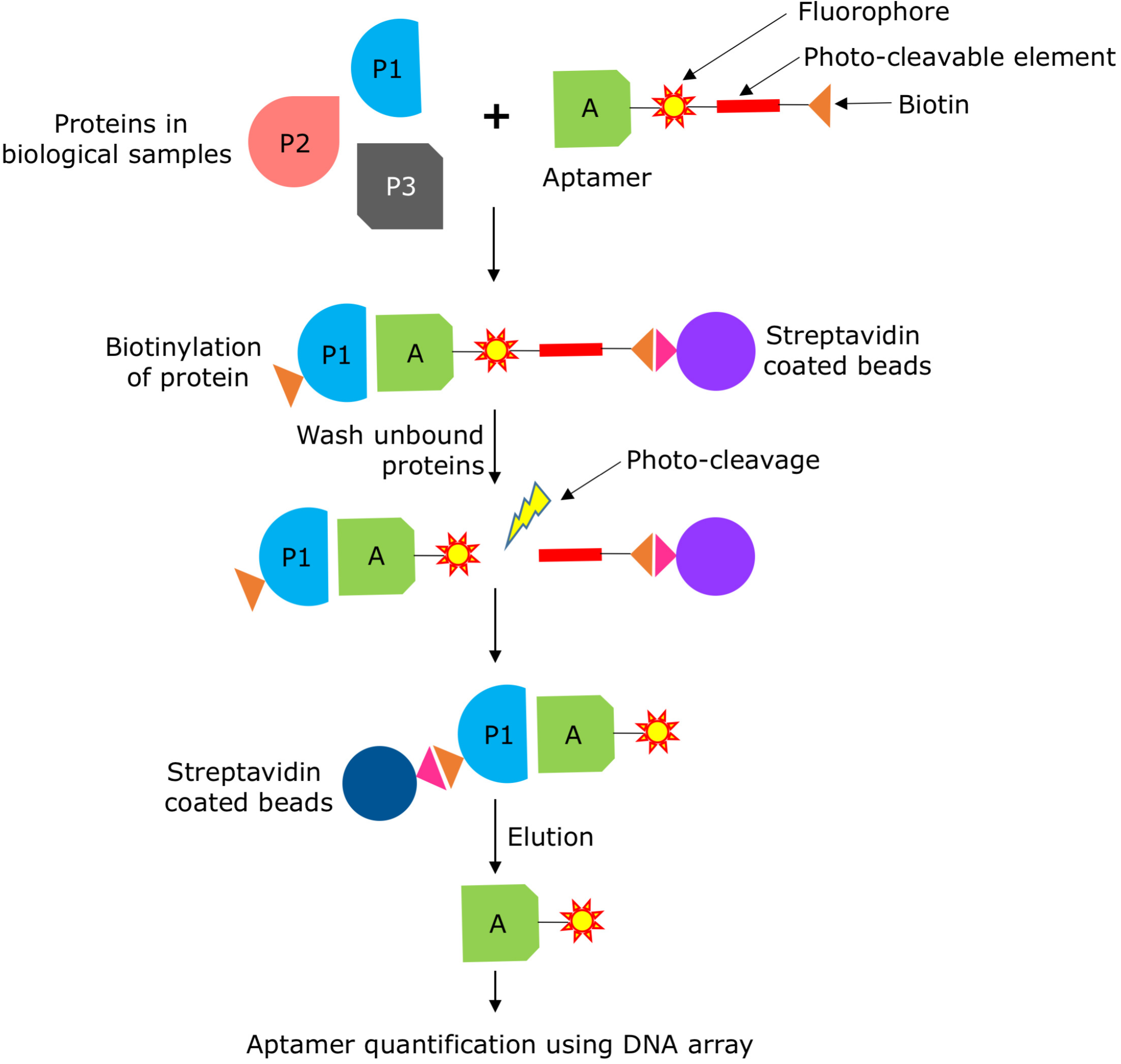
SOMAscan aptamer assay. Proteins in biological samples are mixed with aptamers immobilized on streptavidin beads. The bound proteins are then tagged with NHS-biotin. To prevent non-specific interactions, the beads are washed with an anionic solution. The aptamer-protein complex is then released from the beads by targeting the photo-cleavable linker at the at a peak wavelength of 365 nm. The eluate is again incubated with the streptavidin-coated beads and the unbound aptamers are removed by washing. A final elution is performed and the amount of aptamer-protein is quantified by DNA microarray.

**S. Figure 6.**
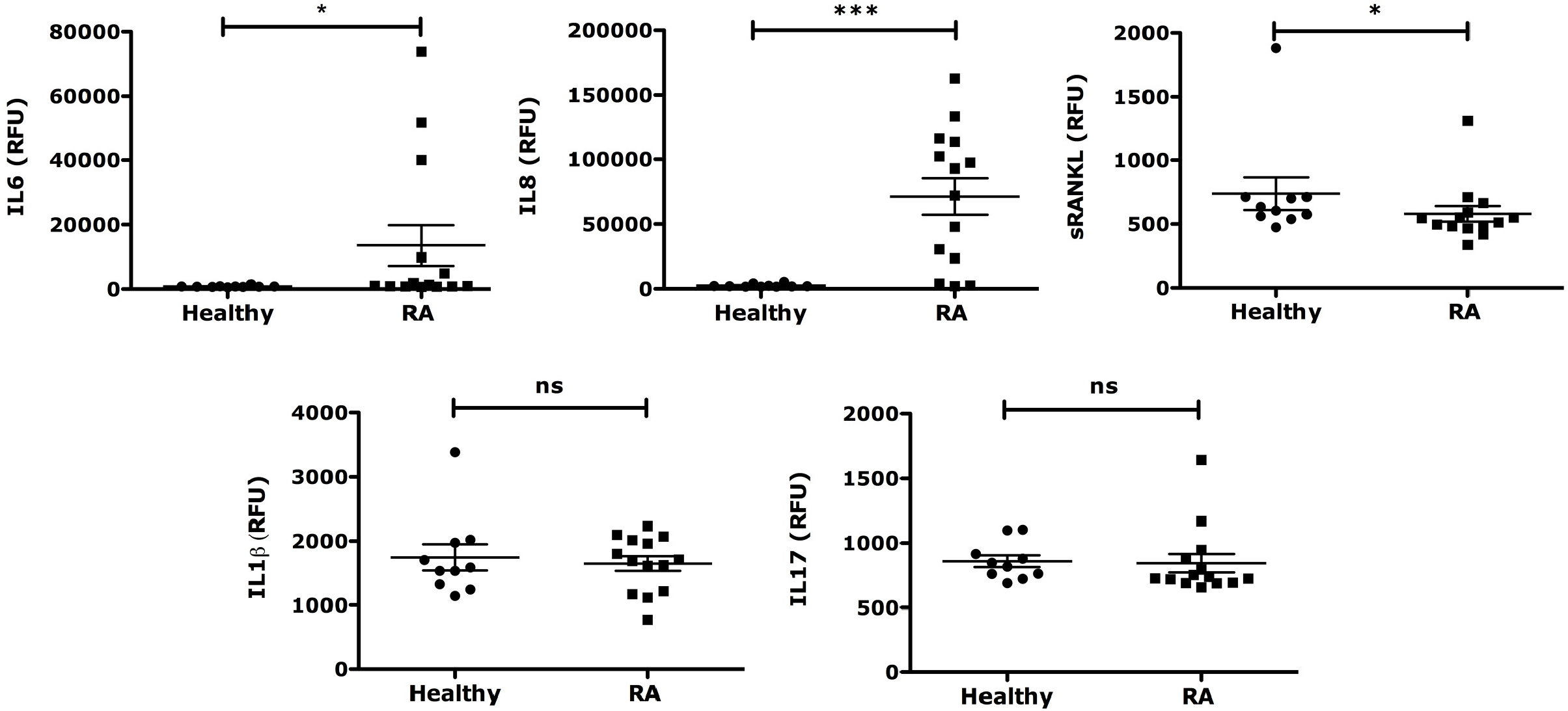
The expression level of key molecules in plasma from RA patients and healthy individuals evaluated by the SOMAscan assay. Higher serum levels of IL6, IL8 and sRANKL were observed in RA patient’s serum, while levels of cytokine IL17 and IL1β are unchanged in comparison to healthy individual’s serum. Statistical analyses were performed using a Mann-Whitney U test.

**S. Figure 7.**
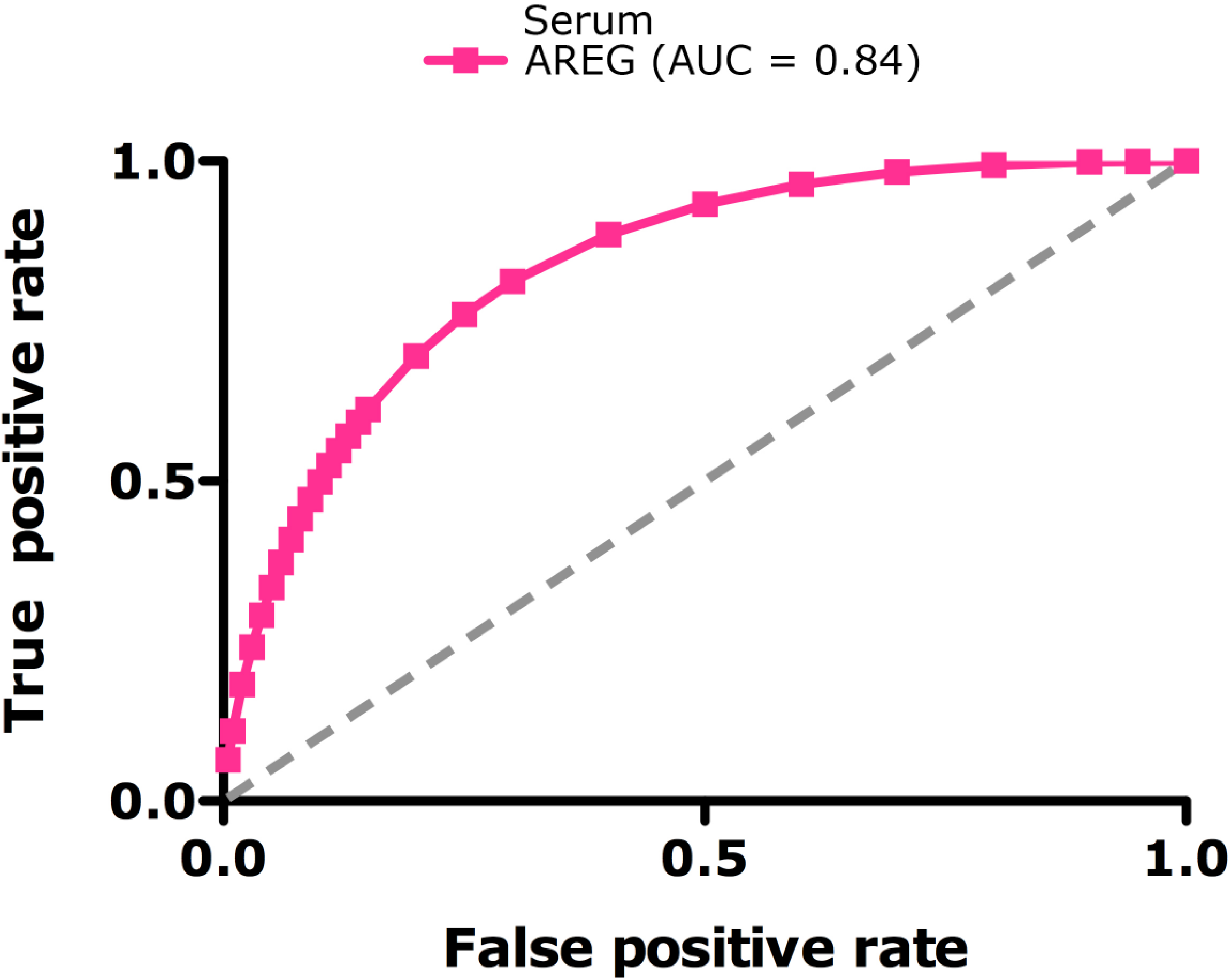
Predictive value of AREG protein in serum for RA. Serum levels of AREG plotted in a receiver operating characteristic curve (ROC) by using the false positive rate and false negative rate of RA occurrence among all serum samples. AUC – Area under the curve

**S. Figure 8.**
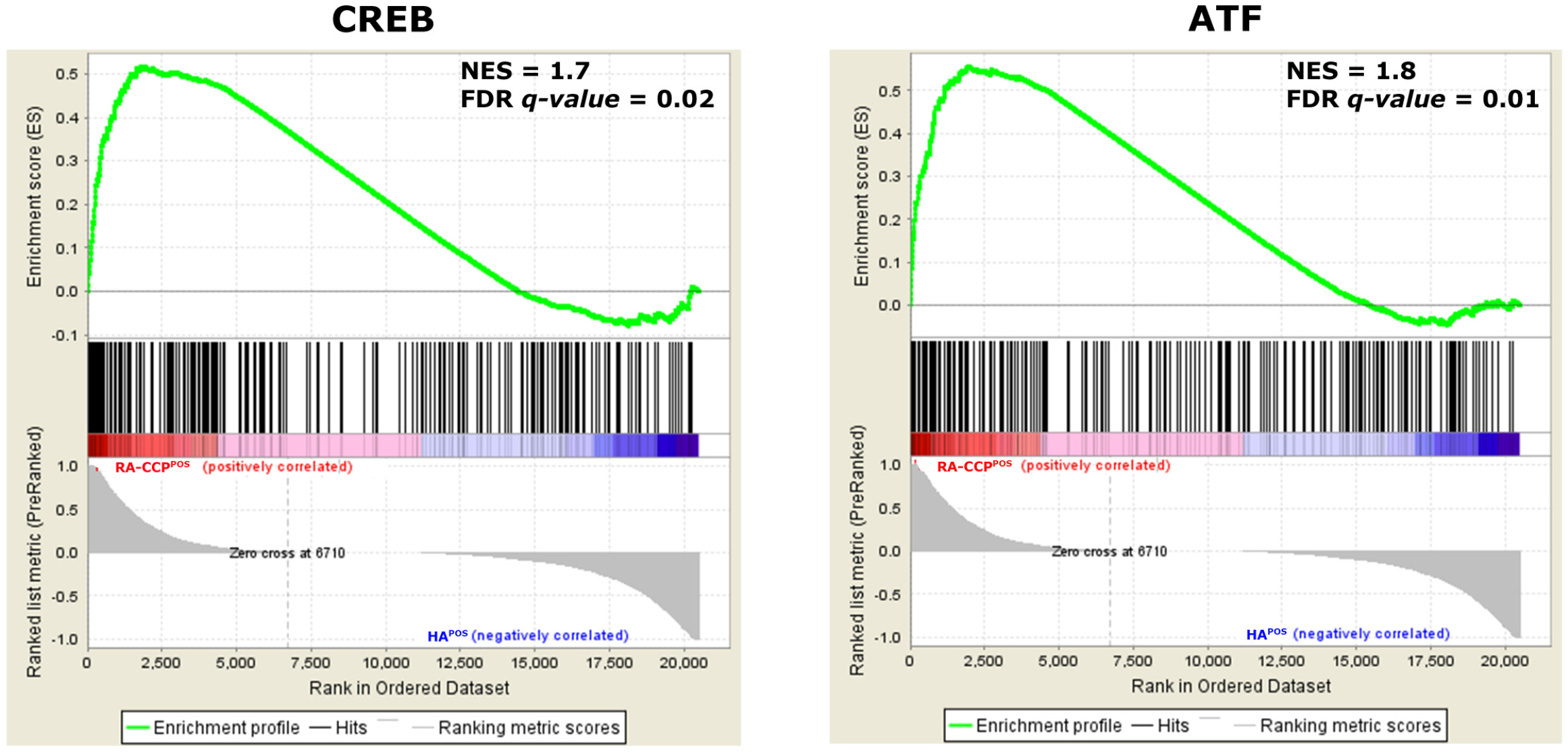
Activation of downstream targets of cAMP. GSEA enrichment plots showing activation of cAMP downstream targets transcription factors, cAMP-response element-binding protein (CREB) and the activating transcription factor (ATF). NES – Normalized enrichment score; FDR – False discovery rate.

**S. Figure 9.**
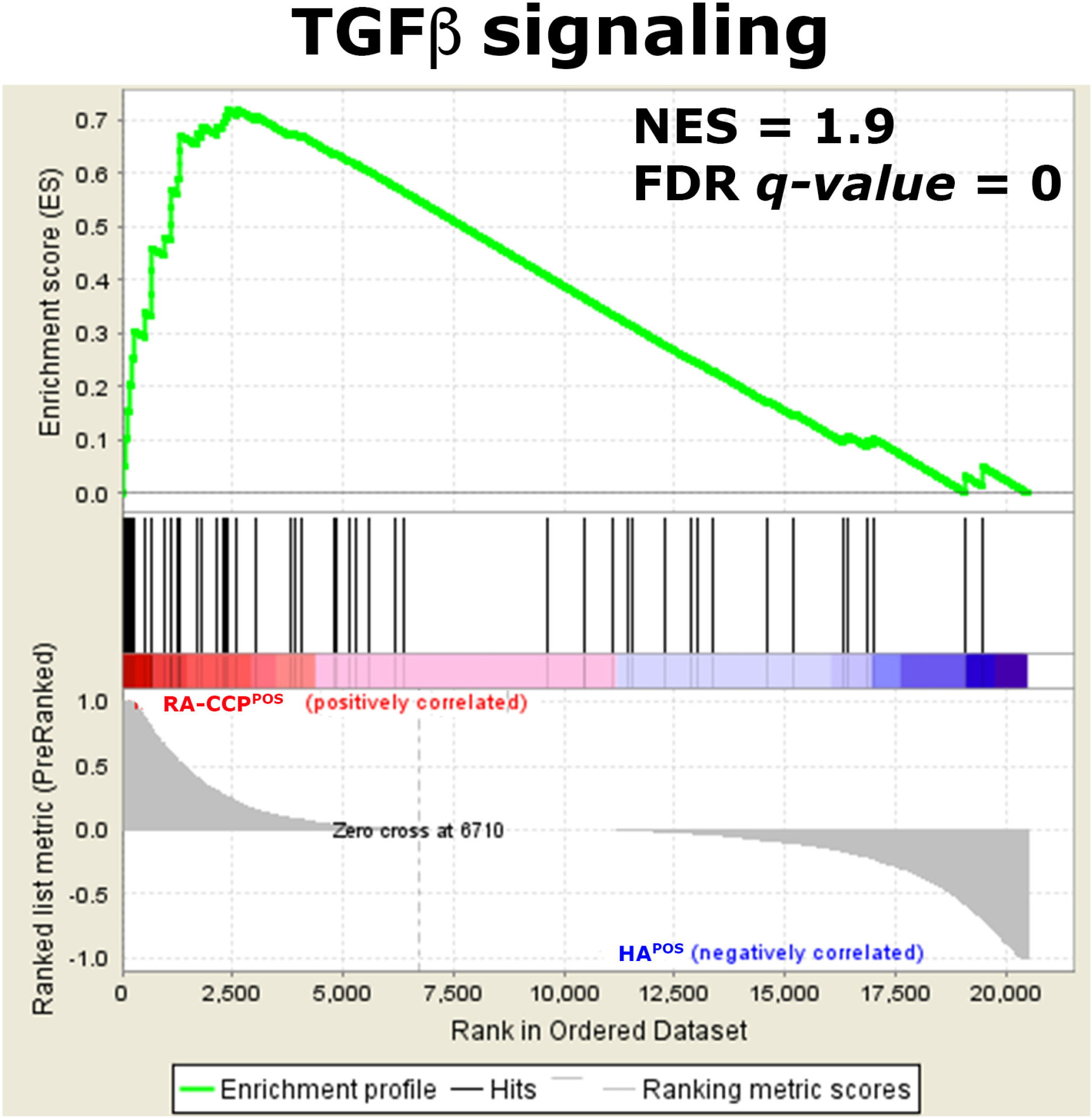
TGFβ signaling is activated in RA. GSEA-derived enrichment plot indicating activation of TGFβ signaling in RA-CCP^POS^ B cells. NES – Normalized enrichment score; FDR – False discovery rate.

